# Benchmarking kinship estimation tools for ancient genomes using pedigree simulations

**DOI:** 10.1101/2023.11.08.566300

**Authors:** Şevval Aktürk, Igor Mapelli, Merve Nur Güler, Kanat Gürün, Büşra Katırcıoğlu, Kıvılcım Başak Vural, Ekin Sağlıcan, Mehmet Çetin, Reyhan Yaka, Elif Sürer, Gözde Atağ, Sevim Seda Çokoğlu, Arda Sevkar, N. Ezgi Altınışık, Dilek Koptekin, Mehmet Somel

## Abstract

There is growing interest in uncovering genetic kinship patterns in past societies using low-coverage paleogenomes. Here, we benchmark four tools for kinship estimation with such data: lcMLkin, NgsRelate, KIN, and READ, which differ in their input, IBD-estimation methods and statistical approaches. We used pedigree and ancient genome sequence simulations to evaluate these tools when only a limited number (1K to 50K) of shared SNPs (with minor allele frequency ≥0.01) are available. The performance of all four tools was comparable using ≥20K SNPs. We found that first-degree related pairs can be accurately classified even with 1K SNPs, with 85% F1 scores using READ and 96% using NgsRelate or lcMLkin. Distinguishing third-degree relatives from unrelated pairs or second-degree relatives was also possible with high accuracy (F1 >90%) with 5K SNPs using NgsRelate and lcMLkin, while READ and KIN showed lower success (69% and 79%, respectively). Meanwhile, noise in population allele frequencies and inbreeding (first cousin mating) led to deviations in kinship coefficients, with different sensitivities across tools. We conclude that using multiple tools in parallel might be an effective approach to achieve robust estimates on ultra-low coverage genomes.

## Background

The use of paleogenomes for inferring genetic kin relations in ancient human populations is growing at an accelerating pace. These studies have unraveled diverse types of social relations of past human societies, from the composition of households [1,2] or burial treatment of mass murder victims [3] to matrilineal [4] or patrilineal traditions studied in graves [5–8]. However, determining kinship degree using single nucleotide polymorphism (SNP) data from low-coverage genomes is fraught with difficulties, mainly arising from data scarcity. The majority of published paleogenomes are below 1x coverage and thus do not allow reliable diploid genotyping, required by popular kinship estimation tools such as KING [9]. Although imputation has recently been shown to produce reliable diploid genotypes using shotgun genomes >0.5x [10,11], a substantial fraction of paleogenomes still do not reach this threshold; e.g., in the AADR repository (v54.1.p1) [12], out of 2041 published shotgun genomes with reported coverage from their original source, 916 (%45) have coverage <0.5x.

A number of solutions finetuned for performance on low coverage ancient DNA (aDNA) data have been published over the last years. These algorithms use pseudohaploid genotypes (e.g. [13]), genotype likelihoods (e.g. [14–16]) or read information (e.g. [17]), instead of diploid calls. These methods also differ in (a) how they normalize the pairwise mismatch values between two genomes to infer the kinship degree, and (b) whether they use method-of-moment estimators or probabilistic approaches. The most widely cited tool, READ [13], compares the rate of average mismatch (P0) between a genome pair with the median (or maximum) P0 of a large enough sample from the same population, assuming this median estimate represents the expected P0 of an unrelated pair. This is similar to the pairwise mismatch rate (PMR) calculation by Kennett and colleagues [4]. Two other commonly used tools, lcMLkin (v2) [14,15] and NgsRelate (v2) [16], use genotype likelihoods and population allele frequency estimates to infer the kinship degree between pairs within a likelihood framework. The TKGWV2 [18] algorithm also uses population allele frequencies within a method-of-moments framework. Finally, the recently published method, KIN [17], uses a likelihood-based framework as well as a Hidden Markov Model (HMM) to infer segments of identity-by-descent (IBD) between pairs of individuals. KIN also uses the average mismatch in a sample for normalizing P0 rates for inferring identity-by-descent (IBD), akin to READ.

Although each of these methods is being widely used by the paleogenomics community, their relative accuracy and performances have not been systematically investigated. One recent exception is a study by Marsh and colleagues [19], who compared these methods using real ancient and modern-day genomic datasets. The authors lacked knowledge of real relationships but studied how consistency among estimates was affected by downsampling high-coverage genomes, reporting that READ, PMR, and TKGWV2 were less affected by low coverage than lcMLkin and NgsRelate. However, this study was limited by the lack of a ground truth set of relationships.

Here, we compare the performances of four commonly used algorithms, lcMLkin, NgsRelate, READ, and KIN, using ancient-like genomic data from pedigree simulations to distinguish close kin (1st-to 3rd-degree relatives) and non-kin. We test the effects of ultra-low coverages (using down to 1000 SNPs per pair), inbreeding, and noise in allele frequency estimates. We chose READ, lcMLkin and NgsRelate as these are among the most widely used algorithms on low-coverage genomes (**Table 1**). Meanwhile, we chose KIN along with NgsRelate as these algorithms are designed to separate genetic correlations due to direct kinship or inbreeding. Importantly, READ and KIN use sample-based normalization, while lcMLkin and NgsRelate use population allele frequencies to infer IBD.

**Table 1:** Different methods and the number of publications using them for kinship estimation. The data was collected by revising literature citing the named articles in Google Scholar (retrieved November 4, 2023) and filtering for publications (including journal publications and preprints but excluding academic theses) that directly used the software (**Supp Table 6**).

| Software | Study | Number of publications<br>using the software |
| --- | --- | --- |
| NgsRelate | Korneliussen & Moltke, 2015 | 47 |
| NgsRelate v2 | Hanghøj et al., 2019 | 57 |
| IcMLkin | Lipatov et al., 2015 | 49 |
| IcMLkin v2 | Žegarac et al., 2021 | 1 |
| READ | Kuhn et al., 2018 | 128 |
| TKGWV2 | Fernandes et al., 2021 | 6 |
| KIN | Popli et al., 2023 | 3 |

## Data Description

We simulated pedigrees with Ped-sim (v1.3) [20] and human sex-specific empirical genetic maps to produce 480 related pairs of individuals, including first-, second-, and third-degree relationships, as well as first- and second-degree relatives with one individual being inbred (**Figure 1**; **Table 2**; Methods). We further collected 29,706 unrelated pairs from the pedigrees. In the simulations, we generated all alternative types of same-degree kinship (e.g., parent-offspring and siblings for first-degree kin) and also all sex constellations because both parameters can change the number of recombinations that separate a pair, and hence the variation in IBD sharing [20,21]. We generated 48 pairs for each of the 8 relationship types (**Table 2**). We created 600 founder genotypes used in the pedigree simulation from the 1000 Genomes Dataset v3 [22] Tuscany (TSI) population SNPs (Methods). The founder genotypes thus involve realistic SNP densities and SNP types, but the founders themselves are artificial and do not carry background relatedness or runs of homozygosity (ROH), which we preferred in order to simplify interpretation. To further render the dataset realistic, we used the Ped-sim generated pedigree genotypes to simulate aDNA-like sequencing data with the gargammel tool for 200,000 SNPs [23]; we then performed the same procedures as applied to standard paleogenome sequencing libraries (Methods). Next, we randomly downsampled the genotypes to a range of shared SNP counts between related and unrelated pairs, from 50K, 20K, 10K, 5K to 1K autosomal SNPs. For each pair and SNP count, we further produced five replicates by randomly downsampling SNP sets. We ran lcMLkin, NgsRelate, READ, and KIN on this data (using perfect information on background allele frequencies with lcMLkin and NgsRelate), and recorded the *θ* (kinship coefficient) and kinship degree assignments. Importantly, we could not run KIN on the sparsest dataset of 1K SNPs, presumably because the algorithm does not converge at such low coverage. We further performed two alternative analyses: (a) we ran NgsRelate after introducing two types of error to background allele frequencies, and (b) we produced genomes with background relatedness using a coalescent simulation and ran NgsRelate and KIN on this dataset (Methods).

**Figure 1:**
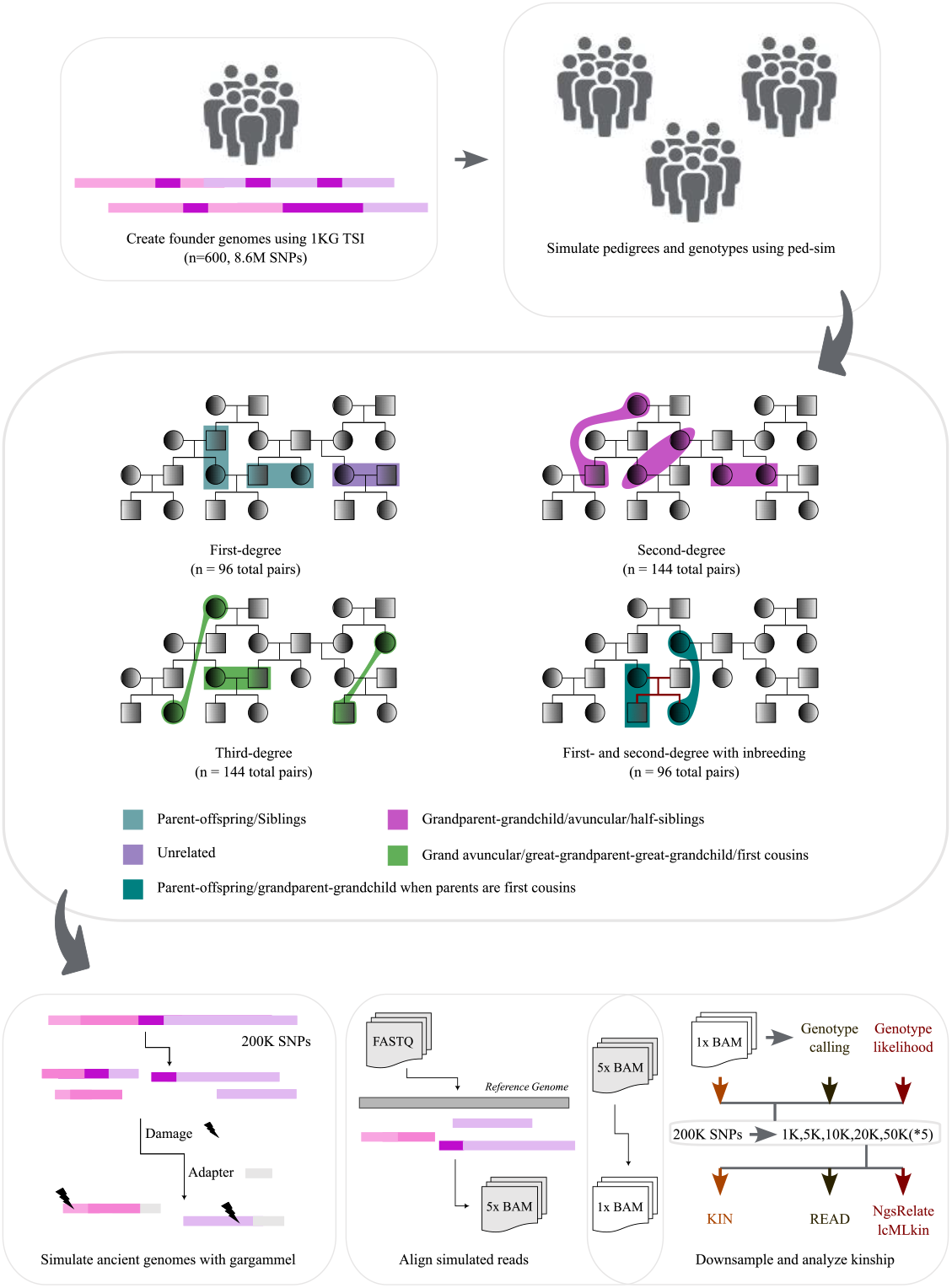
Primary simulations and analysis workflow. For the primary dataset, we created 600 synthetic founder genomes using variant and allele frequency information from the 1000 Genomes Project v3 Tuscany (TSI) sample (Methods). We used these founder genomes to create pedigrees with Ped-sim and human genetic maps, from which we chose sets of related pairs of different types, with n=48 pairs created for each relationship type (2 types for first-degree and 3 types each for second- and third-degree) (Table 2). We also created parent-offspring and grandparent-grandchild pairs where the offspring was the child of first cousins. We subsampled these genotypes to 200K SNPs and created aDNA-like sequencing read data using the gargammel tool around these SNPs. The reads were then aligned to the reference genome to produce 5x BAM files, which were further downsampled to 1x (Methods). We called pseudohaploid genotypes or calculated genotype likelihoods (GL) for the same 200K SNPs and downsampled these to 1K-50K subsets, each SNP counts downsampled randomly 5 times. The genotypes, GL, or BAM files were input into the four kinship estimation tools.

**Table 2:** The relationships used for paleogenomic data simulation. Number of sex combinations: the number of different constellations of the sex of individuals in the same pedigree for each run (e.g. for parent-offspring, this is four depending on whether the parent or the child is female or male). Number of pairs: the number of independently simulated pairs for each type of relationship. “inb”: pairs where inbreeding simulated as the child or grandchild is the offspring of a first-cousin mating (Figure 1).

| Relationship | Degree | Number of sex combinations | Number of individuals | Number of pairs |
| --- | --- | --- | --- | --- |
| Parent-offspring | First | 4 | 72 | 48 |
| Siblings | First | 3 | 96 | 48 |
| Half-siblings | Second | 6 | 96 | 48 |
| Grandparent-grandchild | Second | 4 | 72 | 48 |
| Avuncular | Second | 8 | 96 | 48 |
| First cousins | Third | 10 | 96 | 48 |
| Great-grandparent-great-grandchild | Third | 8 | 72 | 48 |
| Grand avuncular | Third | 16 | 96 | 48 |
| Parent-offspring (inb) | First | 8 | 72 | 48 |
| Grandparent-grandchild (inb) | Second | 4 | 72 | 48 |

## Analyses

### Comparable performances at ≥20K SNPs but weaker results with READ and KIN at lower SNP counts

Both *θ* distributions across all studied pairs and replicates (**Figures 2-4; Figure S1**), the mean *θ* estimates (**Figure 5**), as well as correct kinship degree assignment rates (**Figure 6**) were largely similar among lcMLkin, NgsRelate, READ, and KIN for first-to third-degree relatives and unrelated pairs, using downsampled sets of either 50K or 20K SNPs. As expected, the variance in *θ* tended to be negatively correlated with the SNP count due to random noise, and all *θ* estimates had higher variance between siblings than between parent-offspring due to randomness of recombination (as siblings share ½ of autosomes only on average, while parent-offspring share exactly ½ of their autosomes).

**Figure 2:**
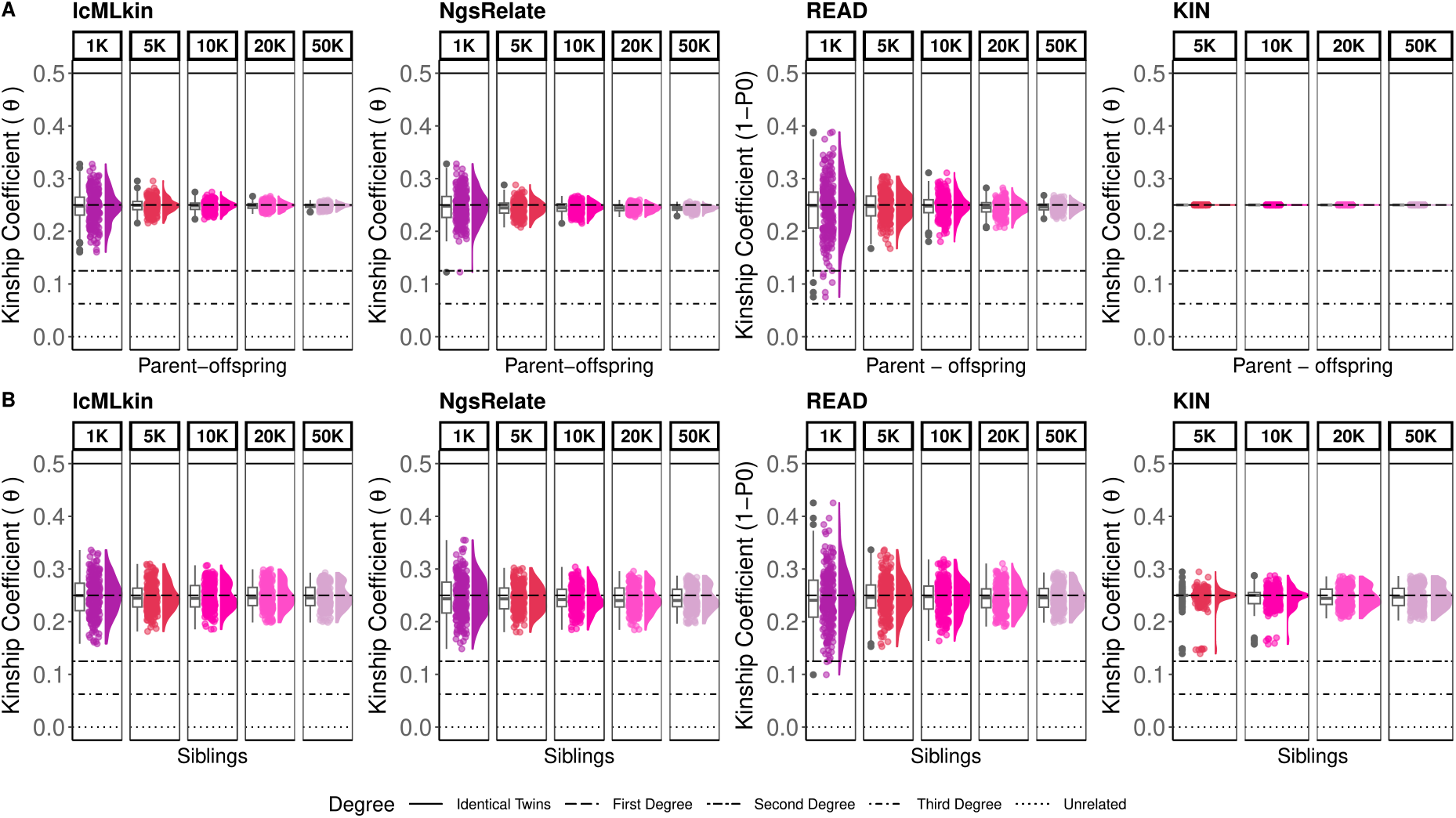
*θ* estimates of simulated first-degree pairs, (A) parent-offspring, and (B) siblings. The points represent the *θ* estimated by lcMLkin, NgsRelate, READ, and KIN for one pair of individuals sharing 1K, 5K, 10K, 20K, or 50K SNPs. KIN results for 1K are missing because the algorithm does not perform at this coverage. For each SNP subset and each relationship type, the total number of simulated pairs is 240. Horizontal lines show the theoretical *θ* values. The boxplots, jitter-added points, and density plots show the distribution of the same sample of 240 points.

**Figure 3:**
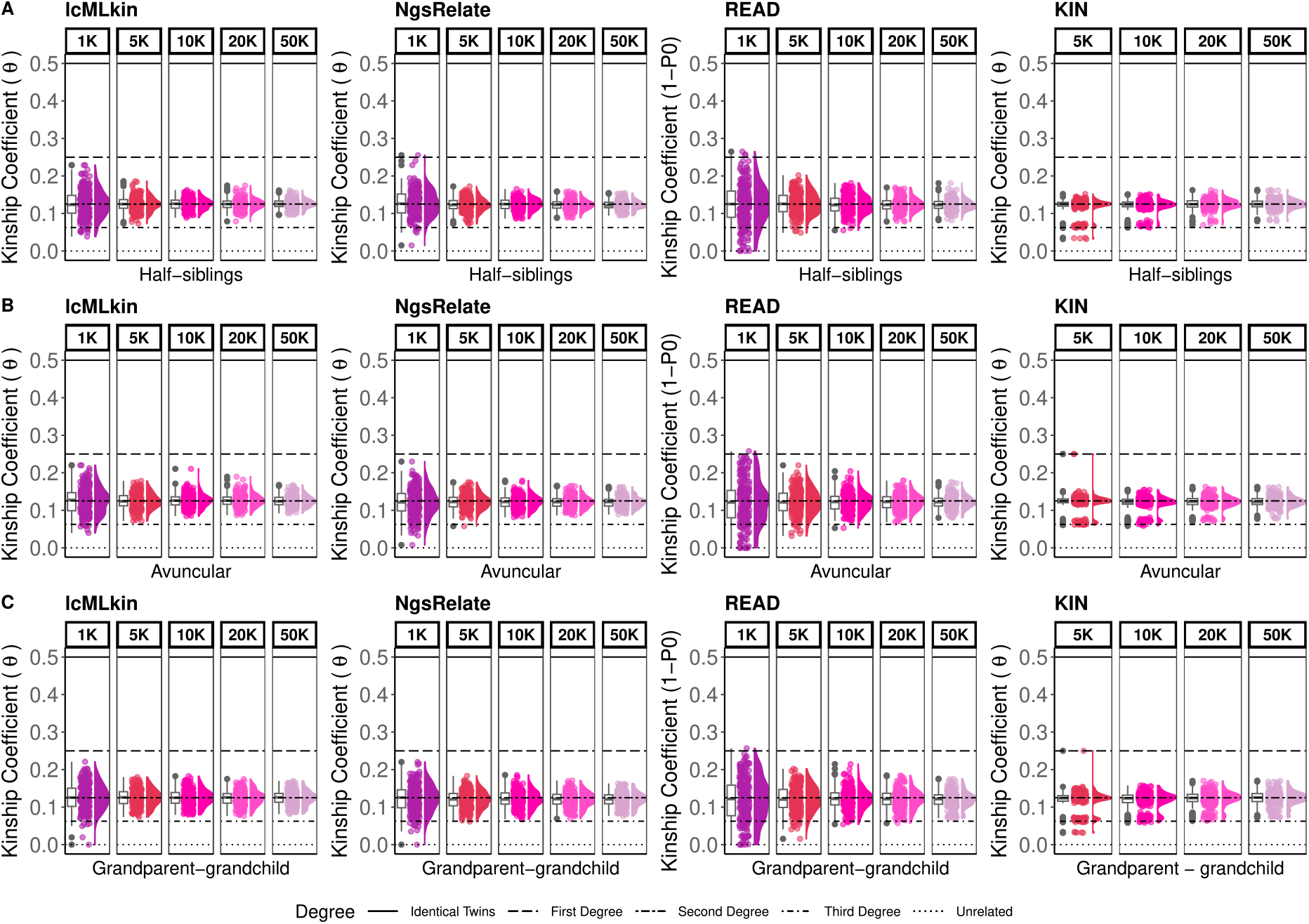
*θ* estimates of simulated second-degree pairs,(A) half-siblings, (B) avuncular, and (C) grandparent-grandchild. The points represent *θ* estimated by lcMLkin, NgsRelate, READ, and KIN for one pair of individuals sharing 1K, 5K, 10K, 20K, or 50K SNPs. KIN results for 1K are missing because the algorithm does not perform at this coverage. For each SNP subset and each relationship type, the total number of simulated pairs is 240. Horizontal lines show the theoretical *θ* values. The boxplots, jitter-added points, and density plots show the distribution of the same sample of 240 points.

**Figure 4:**
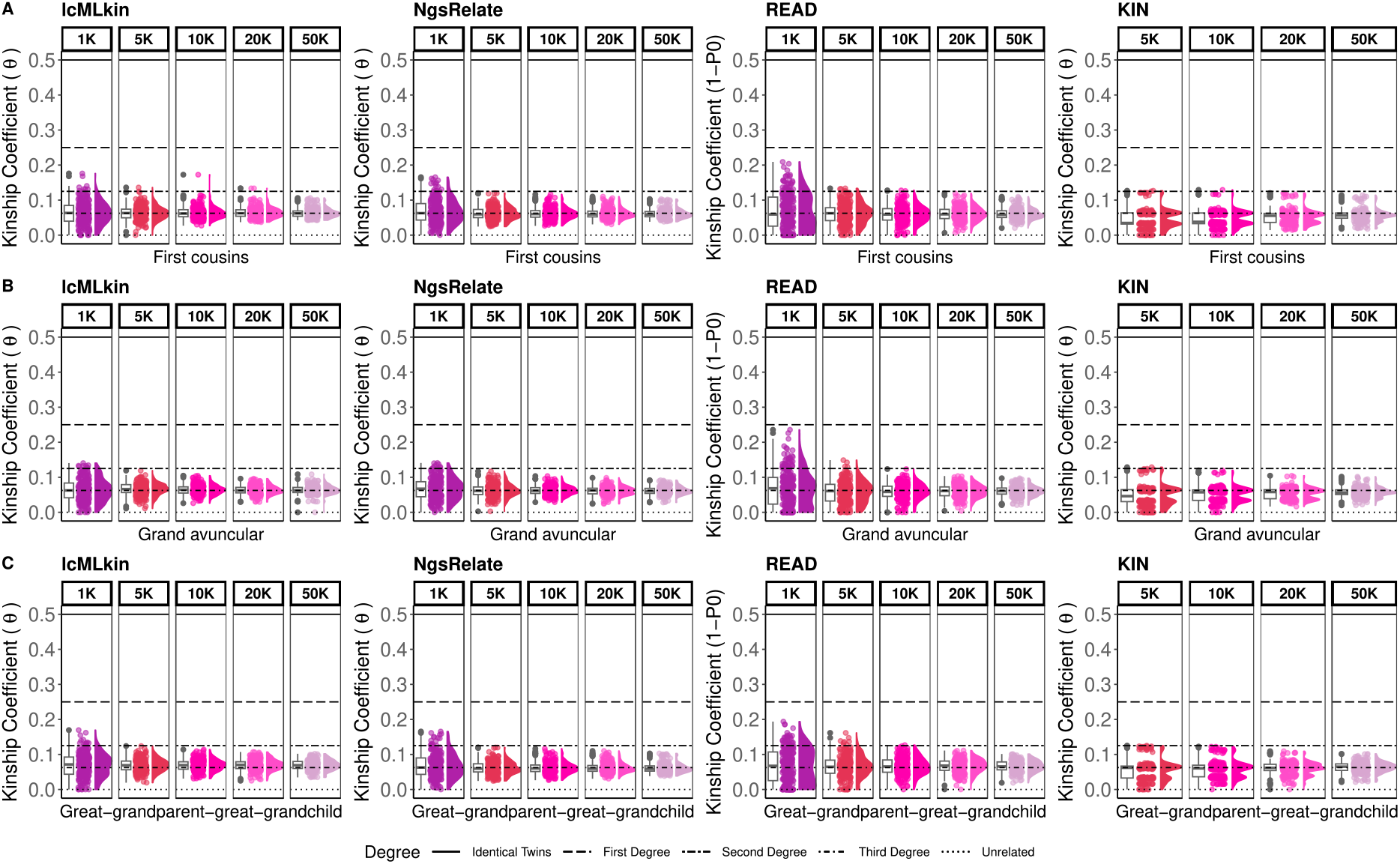
*θ* estimates of simulated third-degree pairs, (A) first cousins, (B) grand avuncular, and (C) great-grandparent-great-grandchild. The points represent *θ* estimated by lcMLkin, NgsRelate, READ, and KIN for one pair of third-degree related individuals sharing 1K, 5K, 10K, 20K, or 50K SNPs. KIN results for 1K are missing because the algorithm does not perform at this coverage. For each SNP subset and each relationship type, the total number of simulated pairs is 240. Horizontal lines show the theoretical *θ* values. The boxplots, jitter-added points, and density plots show the distribution of the same sample of 240 points.

**Figure 5:**
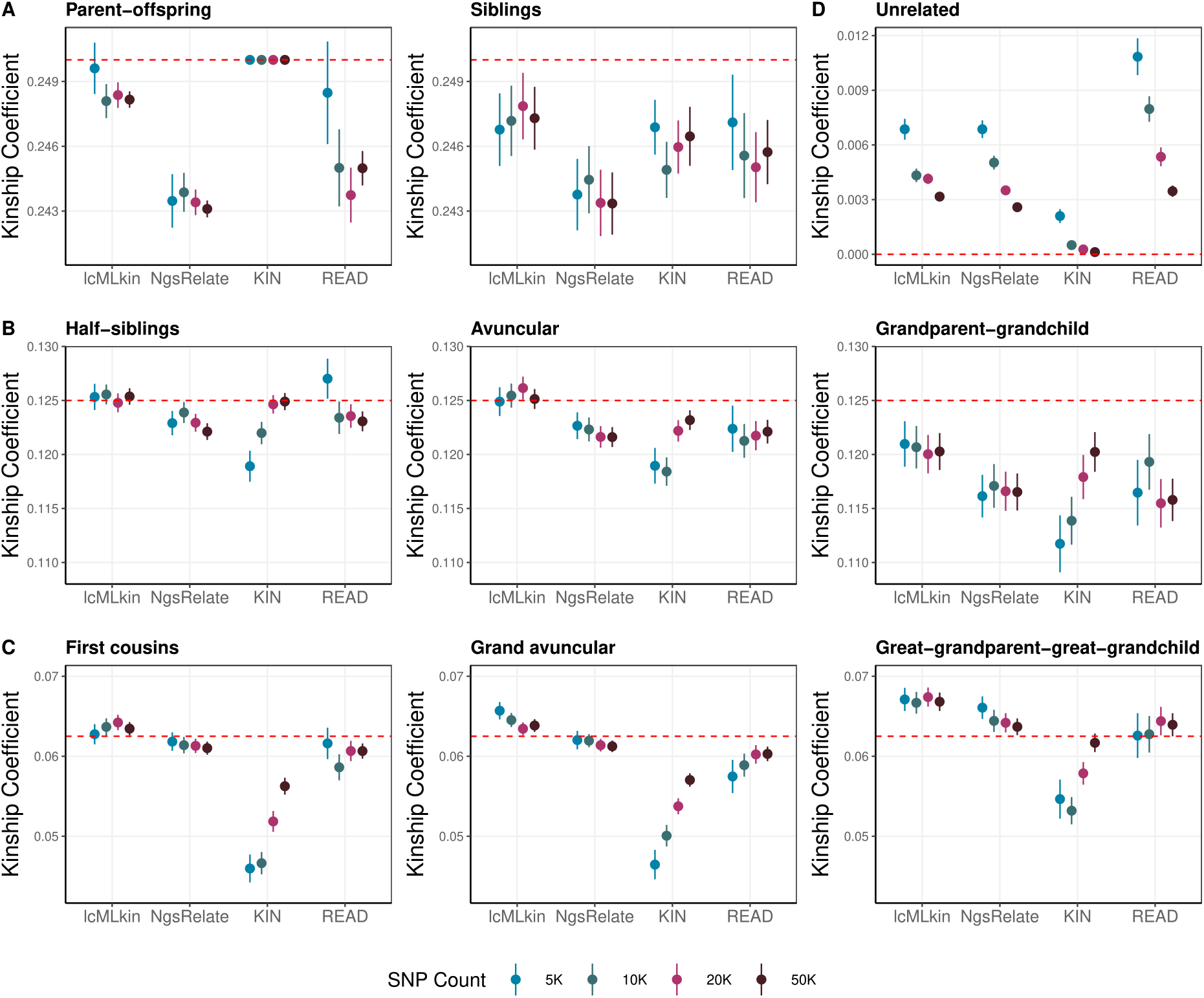
The mean *θ* estimates across different tools and SNP counts for (A) first-degree pairs, (B) second-degree pairs, (C) third-degree pairs, and (D) unrelated pairs, using all pairs (n=48) and replicates (n=5 per pair). Results for each overlapping SNP count are described with distinctive colours. The points show the mean and the vertical lines show +/- one standard error, estimated using all pairs (n=48) and replicates (n=5 per pair). The red dashed line represents the theoretical *θ* value for the corresponding relatedness degree.

**Figure 6:**
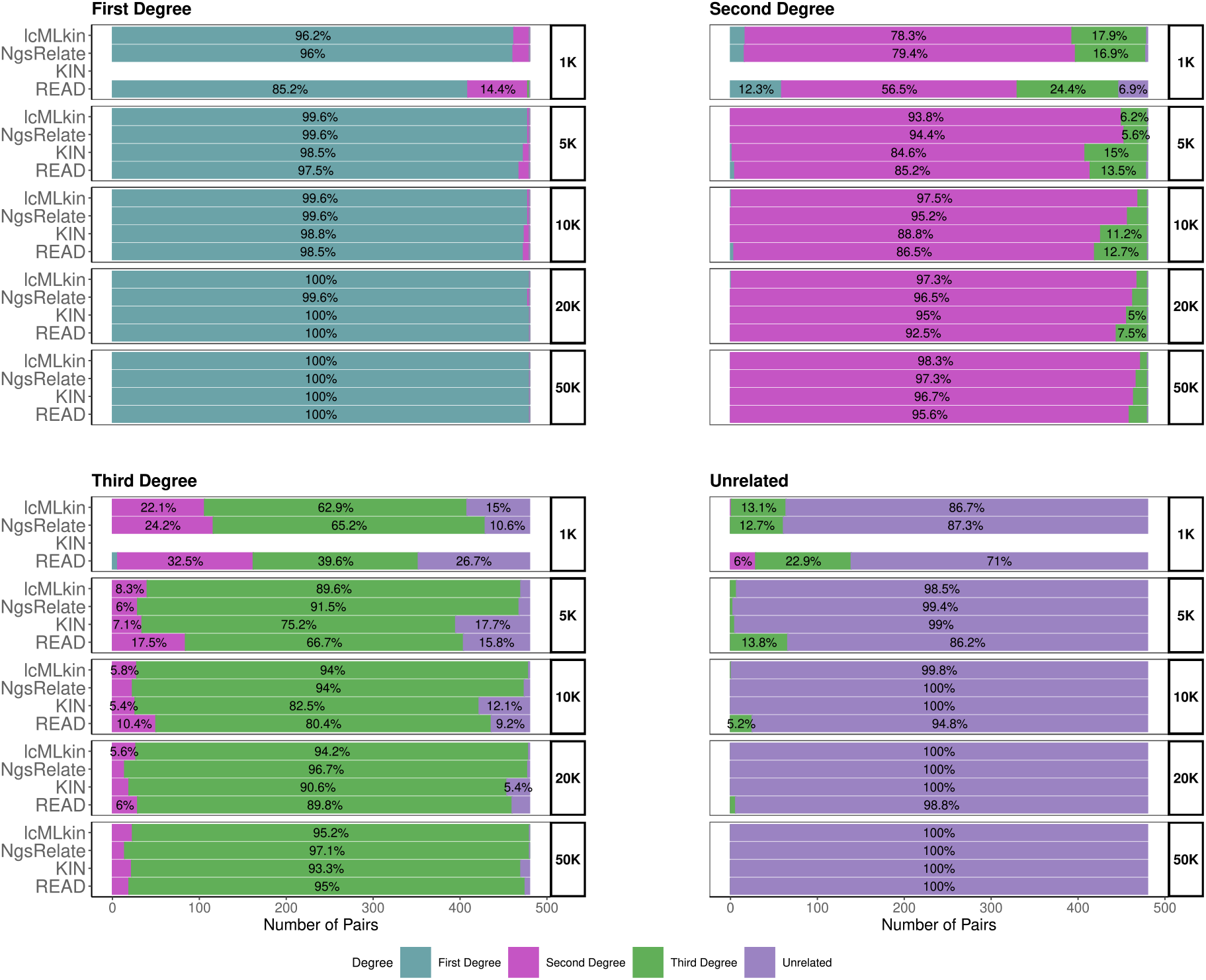
The relative frequency of pairs assigned to first-, second-, and third-degree related and unrelated categories by lcMLkin, NgsRelate, KIN, and READ. The kinship coefficient estimates from these tools were classified using the arithmetic mean of theoretical kinship coefficients. Colors refer to the assigned relatedness degree. The frequencies of pairs assigned to each category are indicated as percentages inside the bars (only for categories with frequency >5%). KIN results for 1K are missing because the algorithm does not perform at this coverage.

We found that identifying first-degree relatives is possible with ≥5K SNPs with all four tools using this dataset with high reliability (≥97.5% correct assignment). Even with 1K SNPs, READ could perform correct first-degree assignments at a frequency of 85.2%, and NgsRelate and lcMLkin at a frequency of >96% (**Figure 6**). Note that KIN did not run with our 1K SNP data as it gave sporadic errors (Methods).

Effectively distinguishing between second-versus third-degree kin, and third-degree kin versus unrelated pairs was more challenging. Still, NgsRelate and lcMLkin reached acceptable performances using down to 1K SNPs and READ and KIN using ≥10K SNPs, with >80% correct assignment.

We also observed a number of systematic differences among the tools. READ performs generally worse than the other three tools with this data in terms of higher variance in *θ* estimates and lower assignment accuracy (**Figures 2-5**). Meanwhile, KIN *θ* distributions have lower variance than the other tools but not improved accuracy, with higher degrees of misassignment than lcMLkin and NgsRelate (**Figure 6**). For instance, using 5K SNPs, the correct assignment of first-degree relatives was 99.6% for both lcMLkin and NgsRelate, compared to 98.5% for KIN and 97.5% for READ. For third-degree relatives, using again 5K SNPs, correct assignment rates were 91.5% for lcMLkin and 89.6% for NgsRelate, in contrast to 75.2% for KIN and 66.7% for READ.

### Bias and variation in *θ* estimates among the four tools

Even though average *θ* estimates are close to expected values under most conditions, leading to correct assignment, slight shifts from expected values can be noticed in **Figures 2-4**. We first inspected these biases among the four tools (Methods). **Figure 5** shows the means of the replicate pairs. One consistent trend was underestimating *θ* in first-degree relationships and grandparent-grandchild pairs and overestimating *θ* among unrelated pairs. Further, KIN diverges from the other tools in displaying the strongest downward bias for related pairs but the least upward bias for unrelated pairs. The observed biases are not strongly correlated with SNP counts, except for KIN estimates. Finally, NgsRelate and lcMLkin appear least biased, but not for all kinship types; e.g. for great-grandparent-great-granchild pairs, READ estimates are closest to expectation. Overall, we find that *θ* estimates from all tools display slight biases, but their level and directions depend on the relationship type and tool (**Figure 5**; **Supp Table 1**). This pattern was also apparent when comparing absolute mean differences from expectation (residuals) using a linear mixed effect model; we tested all 8 kinship types separately, and for each type, at least one pair of software showed significant differences in the magnitude of residuals (at t-test p<0.05) (**Supp Table 2**). These trends, though, appear to have limited impact on classification accuracy: e.g. for siblings, NgsRelate displays the strongest downward bias in average *θ* estimates, but its classification accuracy is higher than both READ and KIN and is on a par with lcMLkin (**Figure 6**). Expectedly, SNP count also had a significant effect on residuals, with larger residuals at lower SNP counts (**Supp Table 2**).

We next studied whether variance among *θ* estimates (as opposed to bias) significantly differs among tools. For this, we ran Levene’s test for variance differences, comparing estimates between the four tools for each relatedness type and SNP count separately (**Supp Table 3**). This revealed significant differences in *θ* variances among the tools, especially with ≤10K SNPs (72/90 of comparisons with p<0.05), which is consistent with their variable classification performance at low coverages (**Figure 6**). The only exceptions were grandparent-grandchild and great-grandparent-great-grandchild pairs, for which variances were similar among tools. The reason for this difference is not obvious but might be related to these kinship types involving fewer observable recombination events than other types [21].

### Higher classification accuracy with NgsRelate and lcMLkin

We next calculated standard accuracy metrics to represent the four tools’ classification performances (**Figure 7**). All tools had high (>98%) F1 accuracy values for first-degree relatives down to 5K SNPs. Even using 1K SNPs, READ had F1 86% while NgsRelate and lcMLkin had F1 96%. Note again that KIN did not perform at this SNP count in our experiments due to sporadic errors (Methods) (**Supp Table 4**).

**Figure 7:**
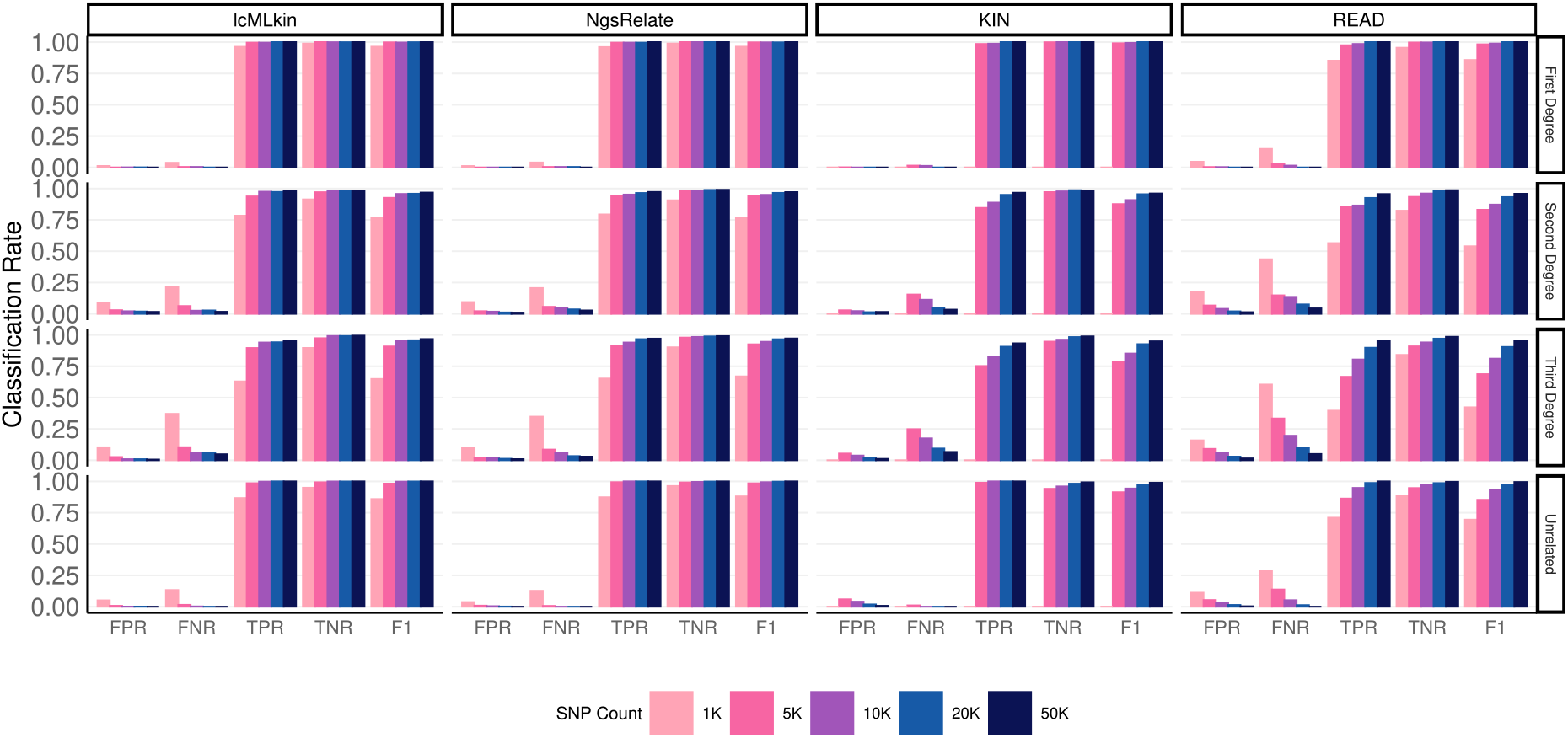
Classification performance of the four tools using the primary dataset. FPR: false positive rate, FNR: false negative rate, TPR: true positive rate, TNR: true negative rate, and F1: accuracy. The classification was performed using n=48 pairs x 5 replicates for each kinship type (n=96 for first-, n=96 for second-, n=96 for third-degree related, n=96 for unrelated), generated using the primary dataset (no inbreeding, perfect background allele frequencies, no background relatedness), and using the arithmetic mean to classify kinship coefficient estimates. Note that we randomly subsampled n=96 pairs for second- and third-degree related categories with each relationship type represented equally (n=32) to ensure balance. The colors represent the count of SNPs shared between individuals. KIN results for 1K are missing because the algorithm does not perform at this coverage.

Beyond first-degree relationships, NgsRelate and lcMLkin performance was superior to those of READ and KIN, especially at low SNP counts. For instance, for second-degree relatives at 5K SNPs, lcMLkin and NgsRelate had F1 values of 93% and 94%, respectively, while READ F1 was only 83%, and that of KIN was 88%, similar to values reported by [17]. This is again consistent with the higher variation of *θ* estimates by READ.

### No major improvement in classification using geometric over arithmetic mean as a threshold

Assignment of pairs to various kinship degrees is traditionally accomplished by using the midpoint between two expected *θ* values (*θ*_1_ and *θ*_2_) as a threshold, i.e. (*θ*_1_ + *θ*_2_)/2 (e.g. [13]). For example, the expected second- and third-degree *θ* values are 0.125 and 0.0625, and thus, the threshold is their arithmetic mean, 0.093, with pairs with *θ* 0.090 assigned to the third-degree kinship class (**Supp Table 5**). Because *θ* and kinship degrees are not linearly correlated (e.g. see **Figure 2**), we asked if the geometric mean 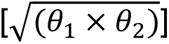, which will be smaller than the arithmetic mean (0.088 in the above case), may provide a more suitable threshold. We ran the classification of the same pairs using the same *θ* estimates from all four tools using the geometric mean as the threshold. We found slightly higher true positive rates using the geometric mean over the arithmetic mean for all categories except third-degree relatives (**Figure S2**). Overall, the differences between the thresholds appear too modest to entail a change in assignment strategy.

### Noise in population allele frequency can lead to over- or underestimation of *θ*

The above results suggest that, at low SNP counts, READ and KIN display lower performance than lcMLkin and NgsRelate. The former pair of tools both use the median mismatch rate in a sample of pairs for normalization, whereas lcMLkin and NgsRelate both use population allele frequency estimates. We reasoned that our use of perfect knowledge of allele frequencies (frequencies used to create the founders) may have favored the performance of lcMLkin and NgsRelate. To study the extent of noise in allele frequency estimates on the latter methods, we performed two additional simulations. Here, we used only NgsRelate, given its highly similar logic and performance with lcMLkin, and only studied 96 first-degree pairs (48 siblings and 48 parent-offspring pairs) for simplicity. First, we introduced random Gaussian noise to the allele frequency estimates with standard deviations of 0.5 and 1 (Methods) (**Figure S3**). As expected, higher random noise led to systematic overestimation of *θ* (>0.25) for all 96 pairs (**Figure 8A-B**; **Figure S4**). This happens because inaccurate background allele frequencies inflate the impact of being identical-by-state (IBS) between any pair.

**Figure 8:**
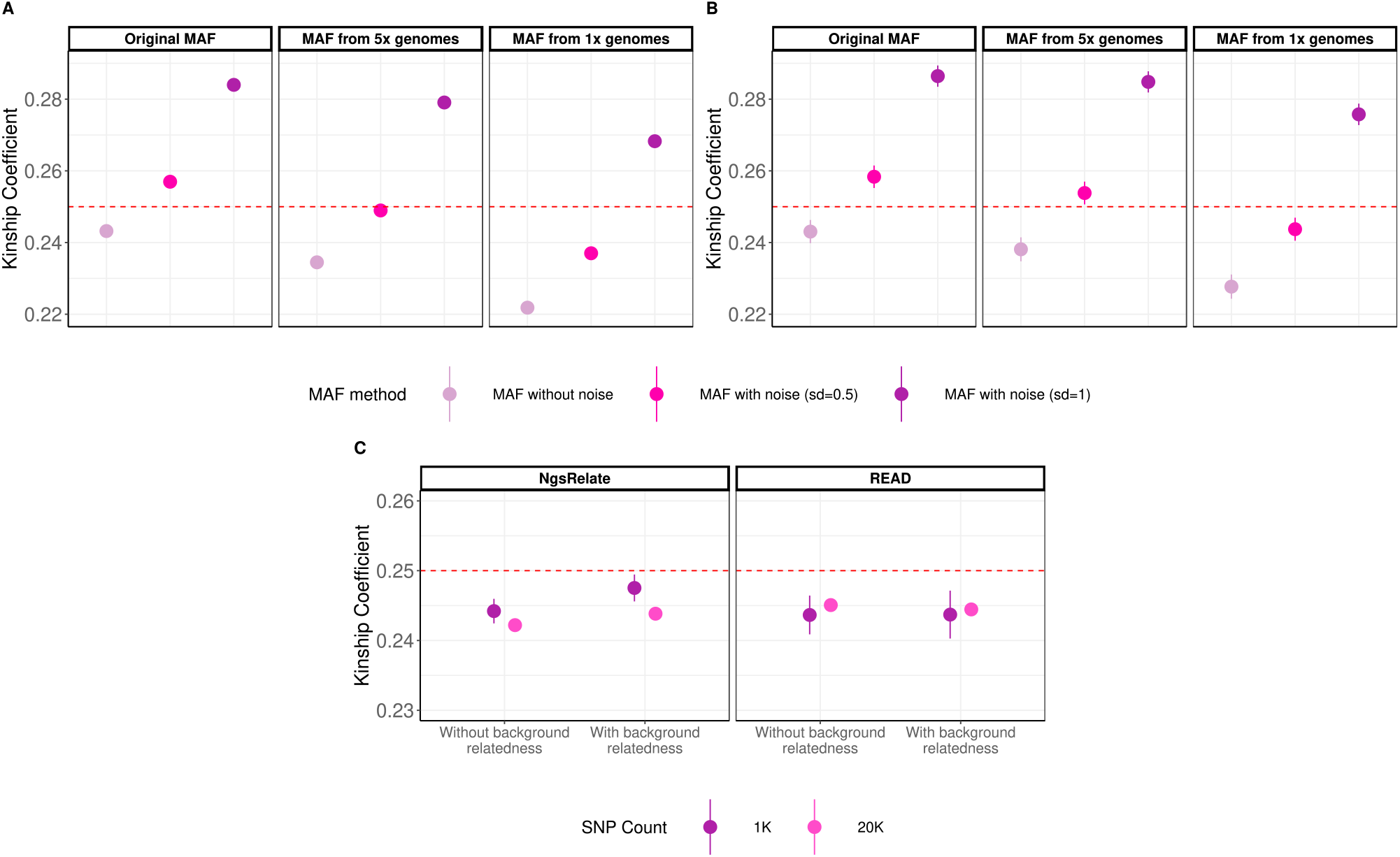
The effects of background allele frequency noise and background relatedness on *θ* estimations. (A) Parent-offspring and (B) sibling *θ* distributions under noise in allele frequencies, calculated using NgsRelate using n=48 pairs each, and all 200K SNPs. “MAF without noise” indicates TSI allele frequencies (perfect information) or MAF from 5x and 1x genomes; “MAF with noise (sd=0.5)” and “MAF with noise (sd=1)” indicate cases where random Gaussian noise is added to allele frequencies; “MAF from 5x genomes” and “MAF from 1x genomes” indicate MAF called using genomes of the indicated coverage (Methods). (C) Parent-offspring *θ* distributions without or with background relatedness using NgsRelate and READ. The points show the mean (n=48 pairs x n=5 replicates) and the vertical lines show +/- one standard error (not visible in panel A) for 1K and 20K SNPs. “Without background relatedness”: the main simulations where synthetic founders were created without background relatedness. “With background relatedness”: simulations where we produced founders using a coalescent simulator and realistic demographic model.

Second, we tested the effect of noise related to imprecise allele frequency estimation. For this, we calculated allele frequencies from 72 simulated genomes of 1x comprising parent-offspring pairs and 96 simulated genomes of 1x comprising sibling pairs, i.e. with limited accuracy (Methods). Intriguingly, this type of noise led to a slight but systematic underestimation of *θ*, with all 48 parent-offspring pairs having *θ* <0.25 and 37/48 sibling pairs having *θ* <0.25 (**Figure 8A-B**; **Figure S4**). The variability among sibling pairs is again likely caused by randomness in recombination. The reason for this underestimation trend could be related to the lower representation of relatively rare variants when estimating allele frequencies from low-coverage genomes (**Figure S3**). Indeed, the underestimation trend was mitigated when using allele frequencies estimated from 5x genomes instead (**Figure 8A-B**; **Figure S4**).

Overall, these results suggest that different sources of noise in population allele frequency estimates can compromise the performance of lcMLkin and NgsRelate. This would also be consistent with the results by Marsh and colleagues [19], who reported low performance of the latter two tools on real genomic datasets.

### Background relatedness has a limited effect on kinship estimates

To investigate whether background relatedness among founders may shift *θ* estimates we produced founder genomes using a coalescent simulator and a demographic model describing European Neolithic ancestry; we then generated a second dataset comprising n=48 parent-offspring pairs from these (Methods). We next ran READ and NgsRelate on 1K and 20K SNP sets and compared the *θ* values with those from the primary dataset with synthetic founders with no background relatedness. We found READ *θ* estimates were practically the same when genomes contained background relatedness, while NgsRelate tended to underestimate *θ* albeit minimally (<0.025) (**Figure 8C**). This suggests that, at least in our simulated scenario of European Neolithic ancestry, the presence of background relatedness among founders might not substantially influence the accuracy or reliability of *θ* estimates produced by READ and NgsRelate using the 1K and 20K SNP sets.

### The effect of inbreeding on *θ* estimates

Inbreeding, either through consanguinity or small population size, can create distal IBD loops between pairs of individuals (**Figure 1**); it will thus increase IBD and elevate *θ* estimates beyond that expected from the proximal relationship. Both past and present human populations are known to vary with respect to average inbreeding levels [24–26]. Among the tools tested here, READ and lcMLkin estimate raw IBD sharing without accounting for inbreeding. NgsRelate estimates the nine Jacquard coefficients (*J*_1-9_) separately and thus could theoretically differentiate between IBD due to proximal loops (*J*_7_ and *J*_8_) versus IBD via distal loops (*J*_3_ and *J*_5_) [16]. KIN, meanwhile, estimates runs of homozygosity (ROH) created by inbreeding in each genome and takes into account ROH-induced IBD when estimating the IBD-sharing level between a pair [17].

We tested the four tools first using parent-offspring simulations, where the parents of the offspring were the first cousins. Average *θ* from READ, lcMLkin, and NgsRelate were 0.27-0.28, as expected (**Figure 9A, Figure S5**). KIN estimates were all 0.25 (except for a single pair using 50K SNPs), suggesting that this algorithm effectively accounts for IBD caused by inbreeding. For NgsRelate, we also calculated a modified *θ* version, 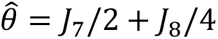, which is expected to reflect proximal IBD sharing without IBD due to distal loops. These 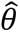 estimates were slightly but systematically lower than what would be expected from proximal loops (∼0.24 using ≥5K SNPs).

**Figure 9:**
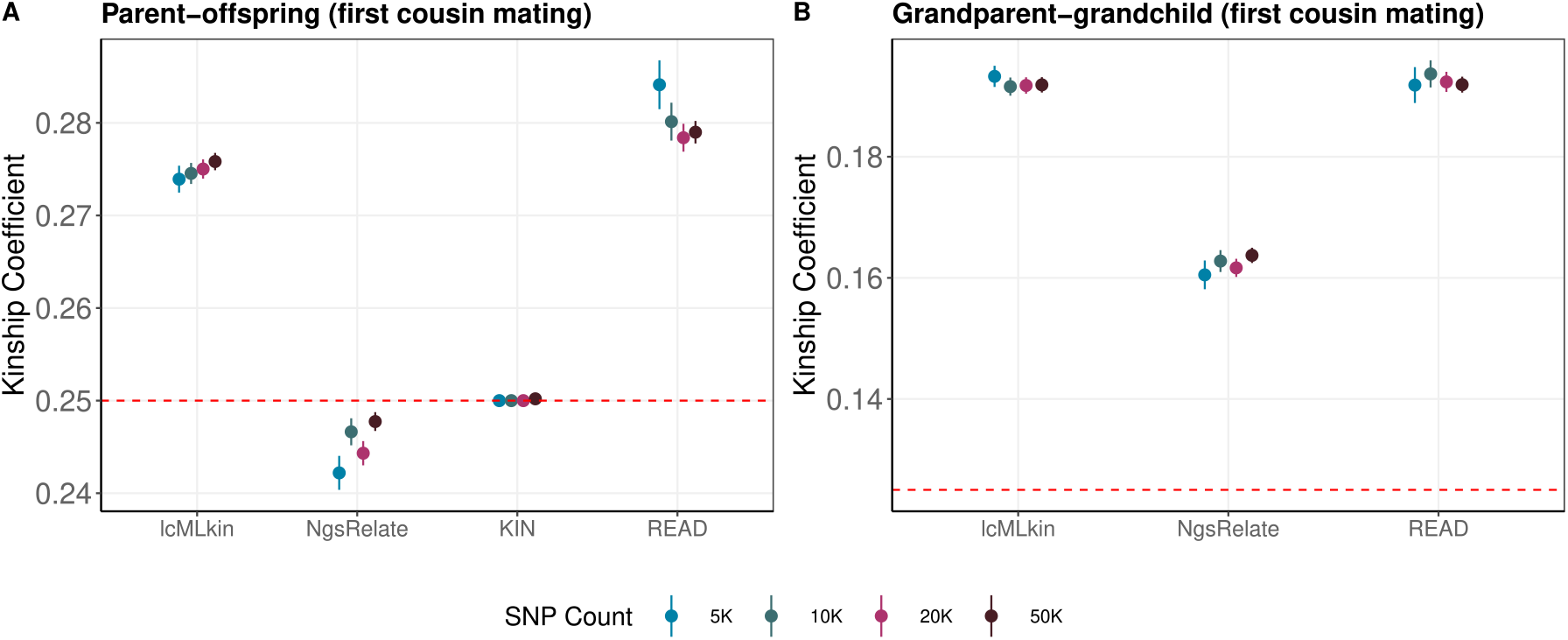
The mean *θ* estimates across different tools and SNP counts for (A) parent-offspring pairs (first cousin mating) and (B) grandparent-grandchild pairs (first cousin mating). Results for each overlapping SNP count are described with distinctive colours. The points show the mean and the vertical lines show +/- one standard error, estimated using all pairs (n=48) and replicates (n=5 per pair). The kinship coefficient from NgsRelate (^*θ*^^) was calculated ignoring the inbreeding-related Jacquard coefficients: ^*θ*^^ = *J*_7_ + *J*_8_/4. The red dashed line represents the theoretical kinship coefficient value for the corresponding relatedness degree. KIN results are missing for grandparent-grandchild results because the algorithm did not perform with this dataset (Methods).

We also simulated grandparent-grandchild pairs, with the grandchild being the offspring of first cousins. Interestingly, KIN gave an error when we ran it with this data (Methods). READ, lcMLkin, and NgsRelate *θ* values were higher than expected from proximal loops (**Figure 9B, Figure S6**). This time, NgsRelate 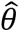 values were also overestimated, but at a lower degree than the above three *θ* estimates.

NgsRelate also estimates individual inbreeding coefficients, *F*, which should be 0.0625 for first cousin mating. The NgsRelate mean *F* estimates for the child were 0.075 for 1K SNPs, but 0.051-0.055 for ≥5K SNPs in the parent-offspring dataset; likewise, mean *F* was 0.068 for 1K SNPs but 0.041-0.048 for ≥5K SNPs in the grandparent-grandchild dataset, suggesting that NgsRelate tends to over- or underestimate *F* in some settings.

## Discussion

Our benchmarking revealed a number of interesting observations on the four tools tested. First, all tools perform well and are consistent with each other down to 20K shared SNPs, even in the separation of third-degree and unrelated pairs (**Figure 6**). This SNP count lower limit corresponds to two genomes each with ∼0.1x coverage genotyped on the 1240k SNP panel [12,27], or each with ∼0.06x genotyped on a 1000 Genomes v3 Africa diversity panel of ∼4.7 million SNPs [28]. Theoretically, this lower limit also applies to comparisons between a 1x genome and a 0.004x genome, using the latter SNP panel.

Nevertheless, we mark that these results reflect upper bounds for performance in real datasets, for a number of reasons:

(a) Our lcMLkin and NgsRelate analyses use perfect information on background allele frequencies, which may be slightly or highly unrealistic in real settings, depending on the dataset.
(b) Our sets of sample pairs used for normalizing mismatch rates, used by READ and KIN, do not include population structure, which would have led to an overestimation of kinship degree as pointed out by Popli and colleagues [17].
(c) Our primary genome simulation dataset lacks background relatedness among the founders, which would be present at variable degrees in real data and could confound estimates of proximal IBD. This involves results from all four tools. Still, our experiment with founders obtained from a realistic demographic model did not create a major shift in *θ* estimates.
(d) We did not include identical genomes or fourth-degree kin in the simulations. This would have lowered accuracy in the classification of first-degree and third-degree categories, respectively.

In our primary simulations, NgsRelate and lcMLkin were found to be more accurate than READ and KIN, with lower false positive and false negative rates, especially when using <20K shared SNPs. The former tools both use genotype likelihoods and population allele frequencies. However, as our trials with noise-added or imperfectly estimated population allele frequencies reveal, this performance might be compromised in real-life applications. In fact, in our own experience, READ results appear highly robust and reproducible compared to those of other tools (e.g. [2,29]).

Another interesting observation was that KIN, which includes inference of both ROH and shared IBD segments using HMMs, did not perform much better than READ in accuracy. We also could not successfully run KIN on 1K SNP datasets and one dataset that included inbreeding. Still, among the three tools tested, KIN is unique in providing likelihoods for kinship degree assignment, as well as separating parent-offspring and sibling pairs.

Overall, our results suggest no single tool may be universally superior in estimating kinship levels with low-coverage genomes. Using multiple tools in parallel and interpreting the results in light of the superiorities and weaknesses of each tool and the particularities of each dataset (e.g. knowledge of allele frequencies, genetic structure within the sample, and the possibility of inbreeding) may be the most prudent approach. Meanwhile, the archaeogenomics community may continue to seek novel and more powerful methods, such as combining the two alternative normalization approaches (population allele frequencies and the median mismatch in a sample) and using haplotype information [30] to calculate more robust kinship coefficients.

## Materials and Methods

### Pedigree Simulations

The goal of this study is to determine how common kinship estimation tools perform on ultra-low coverage ancient genome data. To assess this most effectively, we simulated ancient genome data representing pairs of individuals with known relationships. Briefly, we used pedigree simulation software Ped-sim (v1.3) [20] to produce genotypes from pedigrees of various relationship degrees and types separately, including first-, second-, and third-degree relatedness without inbreeding, as well as first-degree and second-degree relatedness with first-cousin mating. Ped-sim creates individual genotypes based on user-specified pedigrees, using founder individual genotypes and a recombination map (i.e. genetic map) as input.

We created founder genotype data from scratch as follows: We chose autosomal biallelic SNPs with minor allele frequencies (MAF) ≥0.01 from the modern-day Tuscany (TSI) samples (n=112) from the 1000 Genomes Project v3 [22]. For the 8,677,101 such SNPs, we further calculated the alternative allele frequency (AAF) in the TSI. We then created the diploid genotype of each founder by randomly choosing, for each SNP independently, the alternative or reference allele with probability AAF and 1-AAF, respectively, and repeating this twice to create a diploid genotype. Note that this approach eliminates any background relatedness among founders as well as any homozygosity tracts within founder genomes; even though this is not realistic, our choice simplifies the interpretation of the kinship estimation results. We repeated the creation of founder data 12 times (runs), each time producing different sets of founders.

We thus generated 120 unrelated founders (10 for each run, each with n=12) used for first-degree and 240 unrelated founders (20 for each run, each with n=12) for second- and third-degree pedigree simulations each; 600 in total.

We then employed Ped-sim (v1.3) [20] to simulate pedigrees using this founder pool. We used a linearly interpolated sex-specific recombination map [31] with the “*-m*” option and crossover interference model [32] using the “*--intf*” option of Ped-sim. We simulated pedigrees with all possible sex combinations in a relationship (e.g. male-female, female-female, and male-male siblings) by providing “def” files with the “-d” option. We provided Ped-sim the sexes of founder individuals with the “*--sexes*” option. In addition, we used the *“--keep_phase --founder_ids -- fam --miss_rate 0*” parameters for running Ped-sim.

We thus simulated n=72 pedigrees composed of first-degree, n=96 second-degree, and n=96 third-degree related pairs. For instance, for each of the 12 runs generated for first-degree relationships, we chose 6 pedigrees (2 for parent-offspring and 4 for siblings). The founders of each pedigree and simulated individuals from distinct pedigrees were treated as “unrelated”.

From these simulated pedigrees, we chose n=48 pairs for each relationship type (**Table 2**). For instance, for parent-offspring relationships, we chose n=24 parent-offspring trios, n=48 pairs, which resulted in n=24×3=72 unique individuals in total. Overall, the number of unique individuals used for parent-offspring, grandparent-grandchild, and great-grandparent-great-grandchild relationships was n=72 each, while the number of unique individuals used in sibling, half-sibling, first cousin, avuncular, and grand avuncular pedigrees was n=96 each.

For the pedigree simulations with inbreeding, first-degree and second-degree pedigrees (parent-offspring and grandparent-grandchild relationships) were simulated in the presence of first-cousin mating (i.e., the parents of an offspring or a grandchild are first cousins, respectively). For these pedigrees with inbreeding, we also used n=48 pairs for each relationship type (**Table 2**).

### Ancient Sequence Simulation

To create realistic ancient genotypes from this simulated genotype data that contains various types of error inherent in aDNA, we simulated aDNA-like sequencing data and processed this using our standard pipeline for paleogenome sequencing data (see section “Preprocessing of Simulated Ancient Genomes”). Because our aim was to examine kinship estimation at low SNP counts, we sought to speed up these downstream steps by limiting the genotype data to a smaller SNP set. For this, we used an in-house bash script to randomly downsample the 8,677,101 autosomal biallelic SNPs to 200,000 SNPs and used these genotypes for all pairs of simulated individuals. By limiting the number of reads produced, we could significantly reduce the computation time required for alignment.

We next used the gargammel software [23] to simulate aDNA-like Illumina sequencing read data. This ancient read simulator cuts a given FASTA file into variable short lengths mimicking the distribution of read lengths from aDNA libraries, adds post-mortem DNA damage (PMD), adds Illumina adapters to read ends, and finally, introduces sequencing errors and quality scores to produce ancient-looking FASTQ files. To generate input FASTA files for gargammel, for each individual separately (two files for each individual representing either allele), at each SNP position, we inserted alternative alleles according to their genotype into the human reference genome (GRCh37) via the VCFtools “consensus” command [33]. We then cut the FASTA files into 100 bp sequence intervals surrounding each of the 200K SNPs (50 bp on each side) using BEDtools command “get fasta” [34]. For aDNA read size distribution, we used the size distribution file (*sizedist.size*) from gargammel with “*-s*” option, but we removed values higher than 120 bp, resulting in a distribution with a mean of 66.2 bp and a median of 61 bp, ranging between 35 bps and 119 bps. We specified the deamination patterns as “*-damage 0.024, 0.36, 0.009, 0.55*” using the Briggs model parameters [35]. Sequencing errors were introduced using default parameters. We thus generated ancient read data with 5x depth of coverage per individual, without any present-day human or microbial contamination by specifying “*--comp 0,0,1*” option.

### Preprocessing of Simulated Ancient Genomes

We processed the gargammel-simulated read data following the same procedure as applied to ancient genome sequencing libraries in our group and other research teams (e.g., [2,28,29]). Firstly, we removed the adapters from the simulated ancient reads and then merged the paired-end reads [36]. Secondly, the generated single-end ancient reads were mapped to a human reference genome (hs37d5) using the bwa software “*samse*” function (v0.7.15) [37] with the “*-aln*” option, and parameters are set to *“-l 16500*”, “*-n 0.01*” and “*-o 2*”. We eliminated the reads with a minimum of 10% mismatches to the human reference genome. Finally, the remaining reads were trimmed 10 bps from both ends to remove the PMD-related C-to-T and G-to-A substitutions using the bamUtil software with the “*trimBAM*” option [38].

### Genotyping and Downsampling

After Illumina sequencing read simulation and alignment, we randomly downsampled the BAM files of all simulated individuals from 5x to 1x coverage using Picard Tools DownsampleSam (2.25.4) [39]. Because our goal is to study the performance of the kinship coefficient estimation (*θ*) by READ, NgsRelate, lcMLkin, and KIN on low-depth ancient data, most of our analyses involve subsamples of the 1x data (only one read per SNP). We used the 5x data only in testing noise in population allele frequencies.

We next performed pseudo-haploid genotyping from simulated ancient genomes with 1x depth of coverage. Pseudo-haploidization is a regular step in most aDNA genome studies (see Section 1.3.5). This was performed using the SAMtools (v.1.9) “*mpileup*” function [40], followed by running pileupCaller (v1.4.0.5) with the “*--randomHaploid*” parameter [41]. Specifically, to generate text pileup files for all BAM files, we used the random subset of 200K autosomal SNPs that we had selected earlier (see Section 2.1). Mapping quality and base quality filters were set to Phred score >30 in SAMtools (v.1.9) mpileup. Second, the output pileup files were given as input to pileupCaller software to produce pseudo-haploid genotype data by randomly sampling one read and recording its allele at each SNP. Third, the output files were converted to packedped format using ADMIXTOOLS convertf package [42] with parameter “*-p*” and then to transpose ped/fam format using PLINK (v1.9) [43]. Last, we retained only non-missing genotype calls for each pair of individuals using PLINK (v1.9) with the option “*-geno 0*” (note that missing SNPs are removed only for the analysed pair). This reduced the number of SNPs from 200K to an average of 77K for 1x depth of coverage. Missing genotype calls in low-coverage ancient genomes led to a considerable decrease in the number of SNPs.

To explore the lower limits of using ancient genomes for genetic relatedness estimation, we randomly took subsets of 1K, 5K, 10K, 20K, and 50K SNPs shared between each simulated pair. This randomized downsampling was repeated five times for each subset. This allowed us to study how much kinship coefficient estimates vary depending on the set of variants used for the analysis. We note that the term, replicate, used for the downstream analysis refers to this repeated downsampling (n=5).

### Simulations with Background Relatedness

In addition to the primary dataset we generated using synthetic founders from the 1000 Genomes Dataset v3 TSI population (n=112), we created another founder dataset comprising 250 founder individuals with background relatedness. For this, we employed the msprime engine [44,45] in the mode of “*HomSap*” from the stdpopsim library [46,47] to simulate the genetic data of these founder individuals. We utilized the “*HapMapII-GRCh37*” [48] with the “*-g*” option as the recombination map. We simulated the 500 haploid genomes descended from the Linearbandkeramik (LBK) population, which can be described as early European farmers of Anatolian descent [49], of the multi-population model of ancient Eurasia model [50], with the “*-d AncientEurasia-9K19 0 500*” option. Note that this ancestry is supposed to be close to that of the TSI [49]. Subsequently, we transformed the succinct tree sequence output generated by the stdpopsim software into VCF using the tskit library [51] “*vcf*” command with the “*--ploidy 2*” option. We then narrowed our analysis to 200K randomly selected SNP positions through a customized bash script. These selected positions were further used to extract reference bases from the human reference genome (hs37d5) using the “*getfasta*” command of BEDtools (v2.27.1) [34]. We estimated the transition:transversion rate statistics from the 1000 Genomes Dataset v3 TSI population (n=112) to assign alternative alleles to the retrieved reference positions. With this information, we stochastically generated alternative alleles for each position in our dataset, employing a customized R script. This approach was instrumental in replicating genetic variation according to the observed rates within the TSI population, offering a realistic distribution of allele frequencies within our simulated dataset. The rest of the pipeline, comprising pedigree simulation, ancient sequence simulation, preprocessing, genotyping, and downsampling was identical to that used to create our primary dataset.

### Genetic Relatedness Estimation Using READ, NgsRelate, lcMLkin, and KIN

#### READ

READ [13] is a non-parametric genetic relatedness estimation algorithm. READ compares pseudo-haploid genotypes between pairs and calculates the proportion of mismatch positions, i.e., the pairwise mismatch rate (P0), in non-overlapping windows of 1 Mbps. READ then calculates the genome-wide average P0 per pair and normalizes this using a P0 value corresponding to an average unrelated pair. This can be either the mean, maximum, or median (default) of all P0 values in a sample, assuming the average pair is unrelated, or it may be a user-specified value.

We ran READ with pseudo-haploid genotype data of the simulated individual pairs using default parameters. For each of the 8 relationship types, each SNP count, and each random replicate separately, we combined all READ results for all pairs into one set. These sets included both n=48 pairs of a specific relationship type (e.g. siblings) and also unrelated pairs from different pedigrees of this type. The number of unrelated pairs varied between 2468-4458 across relationship types (because some of the pedigrees we produced within the same run included the same founders, we filtered out any pair that shared founders from the “unrelated pairs” category). As these sets were mainly composed of unrelated individuals, we used their median P0 value for normalization (∼0.24), which is also the suggestion of the software developers. The kinship coefficient (*θ*) estimate for each related and unrelated pair was calculated using the formula:

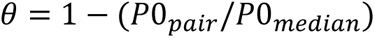

This *θ* estimation approach can yield negative results when a pair shares fewer alleles IBS than the ones of the average unrelated pair [52], suggesting a non-kin relationship. Thus, we set the negative *θ* estimates to 0.

#### NgsRelate

NgsRelate (v2) [16] (hereon NgsRelate) uses maximum likelihood (ML) for estimating genetic relatedness given genotype information and population allele frequencies. NgsRelate further relies on genotype likelihoods (GL) to account for the uncertainty in low-coverage ancient data. NgsRelate uses an expectation-maximization algorithm to estimate nine condensed Jacquard coefficients (J_1_, J_2_,…, J_9_) given GL and population allele frequencies; these coefficients are then used for the direct calculation of kinship:

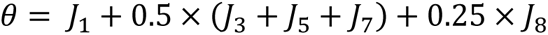

To calculate the GLs for each individual separately from the gargammel-produced BAM files we used the ANGSD program [53] with the “*--gl 2*” option. We limited GL calculation to 200K autosomal SNPs using the “*sites*” parameter for every individual. This left us with 199,095 SNPs passing ANGSD default filters (base quality > 13). The beagle text output file of ANGSD (*--doGlf 2*) was manipulated to generate a GL file containing only two individuals with their shared SNPs. We eliminated pairwise missing SNPs by keeping only sites with GL values not equal to 0.33 for three genotype states (major/major, major/minor, minor/minor) for both individuals with a custom script. Next, we randomly downsampled the shared SNPs between every pair of individuals to 1K, 5K, 1K, 20K, and 50K, five times each, using an in-house bash script. Then, every pair’s GL files with five different SNP subsets were converted to the binary GL file format NgsRelate accepts. The background allele frequency files for corresponding SNPs were prepared using their MAF of the 1000 Genomes TSI sample with n=112 individuals (see below for our NgsRelate trials with alternative background allele frequencies). As the autosomal bi-allelic variants with MAF < 0.01 were excluded from the simulations, the MAF threshold of NgsRelate was set to 0 with the option “*-l* “; this is because the NgsRelate default is 0.05 and we wished to use the same threshold across the software. The output file produced by NgsRelate for each pair includes a *θ* value corresponding to a kinship coefficient estimate. We used this estimated value for subsequent analysis.

#### NgsRelate with alternative background allele frequencies

With NgsRelate, we also conducted trials with alternative background MAF. This analysis was restricted to the two first-degree relatedness categories, parent-offspring (n=48) and siblings (n=48); we reasoned these effects would be consistent across different relatedness types. We ran the ANGSD program with the abovementioned parameters for 200K autosomal SNPs on the BAM files. We processed the resulting GL file to obtain pairwise GL files with no missing SNPs. We then used three alternative background MAF calculations:

(1a) MAFs from the 1000 Genomes TSI population (n=112) as in the original analyses.

(1b) MAFs calculated from gargammel-produced 5x coverage BAM files of the same individuals used in this analysis: 72 individuals comprising the 48 parent-offspring, and 96 individuals comprising the 48 sibling pairs. For this, we ran the ANGSD program with the same parameters on the 5x coverage BAMs and obtained MAFs for both relatedness categories separately.

(1c) MAFs estimated from gargammel-produced 1x coverage BAM files of the same individuals. These were the files used for producing the GL files with ANGSD in the primary analyses.

We also used modified MAFs in three ways:

(2a) No noise.

(2b) Adding a low level of random noise. Here, we introduced random noise to the original MAFs from the TSI while ensuring the resulting values remained within the valid range of 0 to 0.5. For this, we first transformed the MAF values with the logit function: logit(*p*) = log (*p*/(1 − *p*). The purpose of this transformation is to stretch the original allele frequencies to the entire real number space, making them amenable to adding random noise. Then, we generated the noise-added allele frequency values following a Gaussian distribution with a mean based on the logit-transformed MAF values and a standard deviation of 0.5. Then, we applied the expit function (inverse of logit function): *expit*(*p*) = 1/(1 + exp(−*p*)), to the random values to transform them back to the 0 to 1 interval. Lastly, we adjusted the MAF values to ensure they fell within the valid range of 0 to 0.5. This adjustment involved subtracting any values that exceeded 0.5 from 1.

(2c) Adding a high level of random noise. Here, we repeated the same steps as in (2b), but we added Gaussian noise with a standard deviation of 1 (instead of 0.5).

All possible combinations of the three MAF calculations and three noise introductions yielded nine different MAF values (original MAFs and their two different noise-added versions, MAFs calculated from 5x genomes and their two different noise-added versions, and MAFs calculated from 1x genomes and their two different noise-added versions). Then, we ran NgsRelate with the parameters mentioned above for each pair of parent-offspring and sibling categories with these nine different background MAF values.

#### lcMLkin

Another relatedness estimation software using genotype likelihood and population allele frequencies is lcMLkin [14]. Assuming a non-inbred population (unlike NgsRelate) and biallelic loci in linkage equilibrium, lcMLkin estimates the maximum likelihood of Cotterman coefficients also using the Expectation Maximization (EM) algorithm and determines the coefficient of relatedness as *r* = *k*_1_/2 + *k*_2_. Like NgsRelate, the uncertainty in genotype calls of low-coverage NGS data is modeled by summing log-likelihood values of every possible genotype for each site across the genome.

We prepared input VCF files for each pair to run lcMLkin (v2.1) [54] implemented for Python3. For that purpose, we used BCFtools mpileup and call commands [55] to estimate the genotype likelihoods of each individual using BAM files for the 200K SNP set with the mapping and base quality filter parameters “*-q10*” and “*-Q13*”, respectively. These thresholds were selected based on the default filters of ANGSD to estimate GLs for NgsRelate analysis. In this way, we aimed to render the kinship coefficient estimate results from lcMLkin comparable with the estimates from NgsRelate. Besides the VCF files of target samples, lcMLkin requires the genotype data of the selected background population for allele frequency estimation. This genotype data is provided in *PLINK* format (bed/bim/fam) with an argument “*-p*”. We prepared this genotype data using the 200K autosomal SNPs (MAF > 0.01) chosen from the n=112 TSI sample defined earlier. We changed the default allele frequency thresholds integrated into the lcMLkin python script from minimum 0.05 and maximum 0.95 to minimum 0.01 and maximum 0.99. We filtered out missing (non-shared) SNPs from VCF files using an in-house bash script to collect only overlapping SNPs between each simulated pair for the subsequent random downsampling step. After that, we randomly selected 1K, 5K, 10K, 20K, and 50K shared SNPs between pairs of samples, independently five times each, and generated downsampled VCF files using BCFtools view [55] with the “*-R*” parameter. As the LD pruning application of lcMLkin removes closely linked SNPs from the relatedness analysis, we modified the program script such that downsampled SNPs are not pruned by LD. This was done for simplicity to ensure we use the same number of SNPs in each trial and across different software. Also, with ≤50K SNPs across the genome, linkage between neighboring SNPs will be minimal.

The relatedness coefficient (r) is represented with the “*P I_HAT*” estimate in the output files of lcMLkin. We calculated the kinship coefficient value as *θ* = r/2.

#### KIN

KIN [17] has been recently developed to estimate kinship using a Hidden Markov Model-based approach. The properties of KIN that distinguish it from the above-mentioned tools are (i) the ability to differentiate between parent-offspring and sibling pairs, (ii) taking into account inbreeding as inferred from runs of homozygosity (ROH) for relatedness classification, (iii) correcting for contamination. Similar to READ, KIN does not depend on population allele frequencies but estimates P0 in genomic windows directly from read data (BAM files) with a minimum 0.05x depth of coverage. Additionally, it incorporates the probability of window-based ROH tracts in each individual estimated by an *ROH−HMM* model while fitting an IBD sharing pattern of pairs to the predefined relatedness models (unrelated, 5th degree, 4th degree, 3rd degree, 2nd degree, 1st degree and identical) provided by the *KIN−HMM* model. Then, KIN assigns the most likely relationship degree for a pair with the highest likelihood.

As KIN does not work with only two individuals and as we wanted to use one pair at one time to control the shared SNP counts between individuals, we first grouped our BAM files into triplets for each relationship type, including one pair of BAM files to be analyzed and one BAM file of a randomly chosen simulated individual. We determined the read depth of each site at the predefined 200K SNPs for each triplet using SAMtools (v1.9) [40] “*depth*” with the “*-q 30 - Q 30*” options. Then, we removed sites that do not contain at least one read shared between a pair of individuals using a custom bash script since we wanted to keep only shared SNPs for the subsequent analysis.

We thus randomly downsampled remaining sites to 1K, 5K, 10K, 20K, and 50K, independently five times each, for each pair, and gave these downsampled SNP lists as input with “*--bed*” argument to run the KINgaroo algorithm, a python package to generate ROH estimates and input files for KIN. We ran KINgaroo with default parameters without contamination correction (using the “*--cnt 0*” option) and without indexing and sorting of BAM files (using the “*--s 0*” option) for each triplet separately to generate input files necessary for KIN twenty-five times (n=5 SNP counts x n=5 replicates).

Intriguingly, while processing 1K SNP datasets KINgaroo gave sporadic errors, independent of which relationship type was used. Specifically, the algorithm has an “Index Error” (“*IndexError: Can not process input data*”) for several different triplets and replicates with 1K SNPs. Meanwhile, when we ran KINgaroo again with the same triplets but a different set of 1K SNPs without changing any parameter, KINgaroo finished the analysis without error. To further be sure that the problem was related to the usage of 1K SNPs, we continuously ran KINgaroo while using the same or different triplets and different sets of 1K SNPs, but we encountered the same error. We also used the same triplets sharing higher SNP counts (5K, 10K, 20K, and 50K) to run KINgaroo repeatedly; these worked successfully. As we could not understand the reason why the algorithm did not work (possibly could not converge) on some SNP sets, we decided to exclude 1K SNPs and we continued the downstream analysis with higher SNP counts, from 5K to 50K.

We separately collected pairwise mismatch values (P0) of pairs for each relationship type (“*p_all.csv”* file under “*hmm_parameters*” directory created by KINgaroo) and calculated their median P0 values for each SNP count and replicate, corresponding to a P0 value of an average unrelated pair. To apply normalization for kinship estimation with these median values (∼0.24), we manually changed the text files of P0, “*p_0.txt*” created by KINgaroo under the “*hmm_parameters*” directory. We then ran KIN with input files separately for each triplet using default parameters twenty times (n=4 SNP counts x n=5 replicates).

In the grandparent-grandchild relationship with first cousin mating, KIN again did not perform. This time, the program raised an “OS Error” (“*OSError: path/to/directory/likfiles/file1_._file2.csv not found”*). Indeed, we found that KINgaroo had not produced the necessary csv file, although without any warning; the reason for this was again unclear.

The output file of KIN includes the estimates of Jacquard coefficients (k_0_, k_1,_ and k_2_) for each pair analyzed. We calculated the kinship coefficient using these estimates (*θ* = *k*_1_/4 + *k*_2_/2) and used it for the subsequent analysis.

### Classification of kinship coefficient estimates

To systematically test the reliability and robustness of kinship coefficient estimates by lcMLkin, NgsRelate, KIN, and READ on ancient samples, we categorized each simulated pair into one of four relationship categories, i.e., first-, second-, or third-degree related, or unrelated, using their *θ* estimates. Here, we used two assessment criteria. The first criterion we investigated was the arithmetic mean (average) of the theoretical kinship coefficient values. The arithmetic mean of two expected values *θ*_1_ and *θ*_2_ would be (*θ*_1_ + *θ*_2_)/2, i.e. the midpoint of expected kinship coefficient values of two relatedness degrees (**Supp Table 5**). For instance, pairs with 0.1875 > *θ* > 0.09375 would be assigned as second-degree. READ and TKGWV2 also use this mid-point cutoff approach to designate kinship estimates to the appropriate relatedness categories.

The second classification criterion we explored was the geometric mean of theoretical kinship coefficient values. The geometric mean defines the average value of the set of the numbers under study based on their products, and it is always smaller than the arithmetic mean, being closer to the lower value when two values are used. The geometric mean of two expected values *θ*_1_ and *θ*_2_ would be 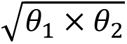. As *θ* values decrease with lower degrees of relatedness in a non-linear fashion (see **Figures 2-4**), we decided to test if using the geometric mean may improve the accuracy of kinship type classification. The cutoffs used are shown in **Supp Table 5**. For the third degree, we determined the threshold using theoretical kinship coefficients of the third-degree related and unrelated pairs, 0.0625 and 0.0, respectively. As zero values cannot be tolerated while calculating the geometric mean, we applied a modified geometric mean for third-degree cutoff using the *splicejam* (v0.0.63.900) package in R [56]. In this way, we derived the third-degree threshold as 0.03078.

### Classification and Accuracy

To compare and contrast the classification performance of the relatedness methods, we created a confusion matrix using either the arithmetic or geometric mean criteria. For this, we used the *confusionMatrix* function of the R *caret* (v3.5) package [57]. Based on the estimated values, this creates a matrix summarizing predictions across a reference or known set of values. In this study, the known values correspond to the relationship categories we simulated: first-, second-, third-degree related, and unrelated.

While producing a confusion matrix and calculating classification metrics in a multi-class scenario like this, it is important to maintain the balance between classes, i.e., an equal number of samples for each class. In our study, the first-degree class includes 96 pairs in total, and it has the lowest number of pairs compared to second (n=144 total, n=48 for each relationship type), third (n=144 total, n=48 for each relationship type), and unrelated (n=29,706) classes (**Table 2**). For this reason, we randomly selected only 96 second- and third-degree related and unrelated pairs using the “*sample*” function of R without replacement. We used the same number of each relationship type for second- and third-degree pairs (n=32 each). After that, we prepared four different datasets for our tools, consisting of classified estimates based on either arithmetic or geometric mean and their actual classes. We separately applied the confusion matrix function to the datasets for each shared SNP count (1K, 5K, 10K, 20K, and 50K).

The metrics we used for benchmarking each of the four tools were the true positive rate (TPR), true negative rate (TNR), false positive rate (FPR), false negative rate (FNR), precision, and the F-score (F1). To understand how often the four software correctly identified the estimates, we also determined the relative frequency of both true and false predictions for each class and SNP count. Additionally, we categorized the false predictions according to their inferred classes using the same confusion matrix again.

### Statistical Tests on Kinship Coefficient Estimates

#### Linear Mixed Effect Model

We used a linear mixed effect model (random-effect model or multi-level model) to study the effect of software choice and SNP count on *θ* estimates for each relationship type. The fixed effects were (a) the type of genetic relationship estimation tools we used, i.e., READ, NgsRelate, KIN, and lcMLkin, and (b) SNP counts shared between simulated individuals (5K, 10K, 20K, and 50K). Here, 1K SNPs were not included because KIN did not perform with this SNP count (see above). The pair of individuals used was included as a random effect. The *θ* estimates were the response variable.

We used the lmer function in the R *lmerTest* package [58] with the R code: *lmer*(*θ*∼*Software* + *SNP*_*Count*_ + (1|*pairs*)). We repeated the analysis with each relationship type separately. We used the R base function “*summary*” on the lmer object to visualize p-values of pairwise mean *θ* difference among software and SNP counts, using lcMLkin and 50K SNPs as the baseline. To ensure data independence, if multiple pairs included the same individual (which happened among parent-offspring, grandparent-grandchild, and great– grandparent–great–grandchild pairs), we chose only one of the pairs, so that our data did not include the same individual in multiple pairs. In this way, we kept only 24 pairs for these three relatedness types.

Additionally, we applied the same linear mixed effect model but this time using as a response variable the absolute residuals, i.e., the absolute differences between the *θ* estimate of a pair and theoretical *θ* value, *AMD* = |*θ*_*expected*_ − *θ*|. This way, we investigated the possible deviations from the theoretical values while accounting for the variances between pairs.

#### Levene’s test

We performed Levene’s test to explore the homogeneity of variances between the kinship coefficient estimates of the tools using the “*leveneTest*” function in the R “*car*” package [59]. We first divided the estimates from READ, NgsRelate, lcMLkin, and KIN into groups based on SNP counts and replicates. Then, we applied Levene’s test separately to each group using their kinship coefficient estimates.

## Authors’ Contributions

Ş.A, M.N.G. and M.S. designed the study. Ş.A., I.M. and M.N.G. produced the data with the support of K.B.V.. Ş.A., M.N.G. and B.K. analyzed data assisted by I.M., K.G., K.B.V., E.S., M.Ç., R.Y., E.S., G.A., S.S.Ç., A.S., N.E.A., D.K., M.S. Ş.A., M.N.G. and M.S. wrote the manuscript with contributions from all authors.

## Data Availability

Simulated genotypes and BAM files were deposited at Zenodo at doi:10.5281/zenodo.10070958, doi:10.5281/zenodo.10079685, and doi:10.5281/zenodo.10079625.

## Acknowledgements

We thank all members of the METU Biological Science CompEvo and of the Hacettepe Human_G groups, Torsten Günther, Gülşah Merve Kılınç, Aybar Can Acar and Burçak Otlu for discussions, Divyaratan Popli and Douaa Zakaria for help.

