## Supplementary Information for "Benchmarking kinship estimation tools for ancient genomes using pedigree simulations"

**Supplementary Table Legends**

**Supp Table 1:** Linear mixed-effects model results on $\theta$ estimates as response variable, pair as random effect, and SNP count and software as fixed effects. Each tab shows results run separately for PO: parent-offspring, S: sibling, HS: half-sibling, AV: avuncular, GPC: grandparent-grandchild, FC: first cousin, GAV: grand avuncular, GGPC: great-grandparent-great-grandchild relationships. Each relationship type was tested using n=48 pairs except PO, GPC and GGPC types tested using n=24 pairs (Methods). In each tab, the regression table on the top shows the estimate, the standard error, the t-statistic, and the t-test p-value, using lcMLkin and 50K SNPs as baseline. In each tab, the ANOVA results at the bottom show the overall p-values for "*software*" and "*SNP count*". We did not include 1K SNPs as KIN did not perform on this dataset.

**Supp Table 2:** Linear mixed-effects model results on absolute mean difference between observed and expected $\theta$ estimates (absolute residuals) as response variable, pair as random effect, and SNP count and software as fixed effects. Each tab shows results run separately for PO: parent-offspring, S: sibling, HS: half-sibling, AV: avuncular, GPC: grandparent-grandchild, FC: first cousin, GAV: grand avuncular, GGPC: great-grandparent-great-grandchild relationships. Each relationship type was tested using n=48 pairs except PO, GPC, and GGPC types tested using n=24 pairs (Methods). In each tab, the regression table on the top shows the estimate, the standard error, the t-statistic, and the t-test p-value, using lcMLkin and 50K SNPs as a baseline. In each tab, the ANOVA results at the bottom show the overall p-values for "*software*" and "*SNP count*". We did not include 1K SNPs as KIN did not perform on this dataset.

**Supp Table 3:** Levene's test results for homogeneity of variances between $\theta$ estimates of the four tools, conducted separately on each SNP count and relatedness category. We did not include 1K SNPs as KIN did not perform on this dataset. Each relationship type was tested using n=48 pairs except PO, GPC, and GGPC types tested using n=24 pairs (Methods).

**Supp Table 4:** Classification metric results for each tool (TPR, TNR, FPR, FNR, recall, precision, and F1) for each SNP count and relationship degree.

**Supp Table 5:** Theoretical kinship coefficient values and classification cutoffs. The cutoffs are calculated using arithmetic or geometric mean of theoretical values for identical/twins, first-, second-, and third-degree, and unrelated classes.

**Supp Table 6:** The list of publications that directly used the software benchmarked in this study. The data was collected by revising literature citing the named articles in Google Scholar (retrieved November 4, 2023) and filtering for publications (including journal publications and preprints but excluding academic theses) that used the software.

### Supplementary Figures

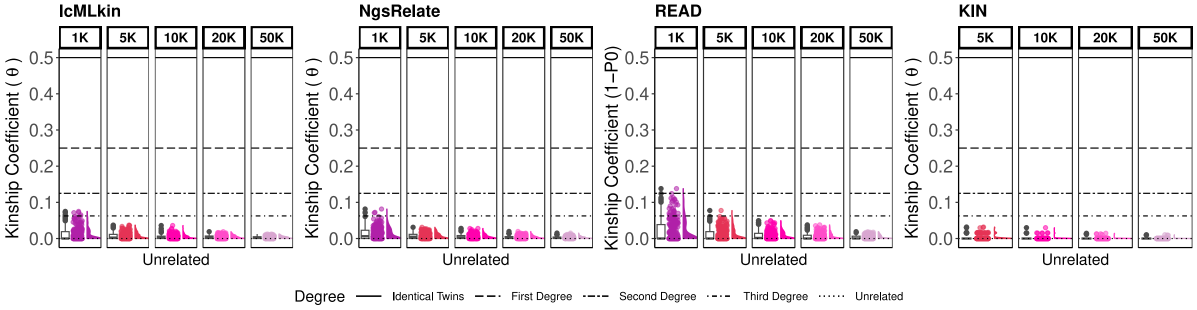


**Figure S1:** Kinship coefficient estimates of simulated 48 (randomly chosen out of 29,706) unrelated pairs across 5 replicates for subsets of 1K, 5K, 10K, 20K, and 50K SNPs. The points represent the kinship coefficient estimated by lcMLkin, NgsRelate, READ, and KIN for one pair of individuals sharing 1K, 5K, 10K, 20K, or 50K SNPs. For each SNP subset, the total number of simulated pairs is 48x5=240. Horizontal lines show the theoretical kinship coefficients. The boxplots, jitter-added points, and density plots show the distribution of the same sample of 240 points. KIN results for 1K are missing because the algorithm does not perform at this coverage.

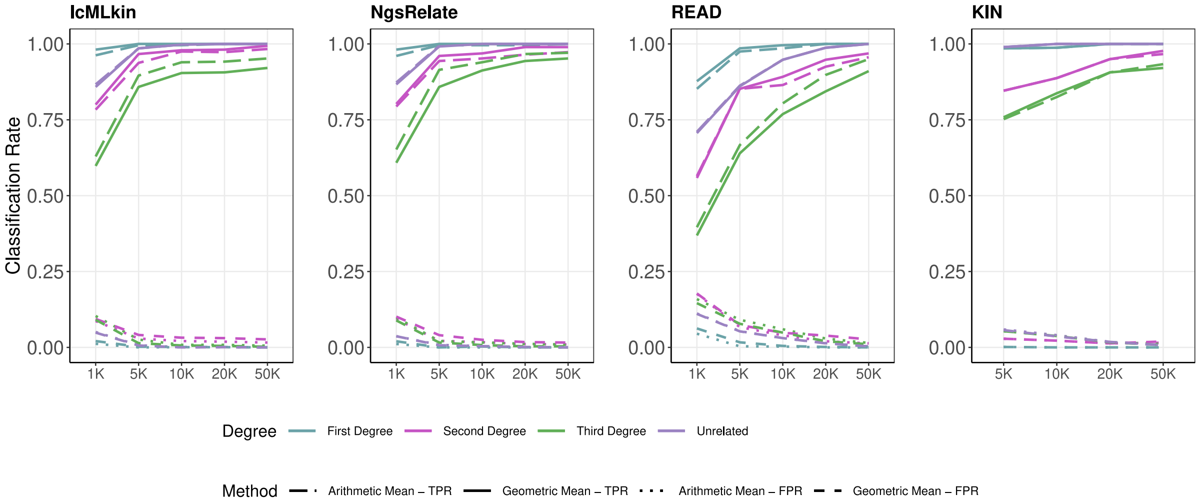


**Figure S2:** True positive rate (TPR, or sensitivity) and false positive rates (FPR, i.e., $1 - specificity$) for lcMLkin, NgsRelate, READ, and KIN using two different classification methods, arithmetic and geometric mean. Colors refer to the relatedness degree, and line types represent the classification method and metric. KIN results for 1K are missing because the algorithm does not perform at this coverage.

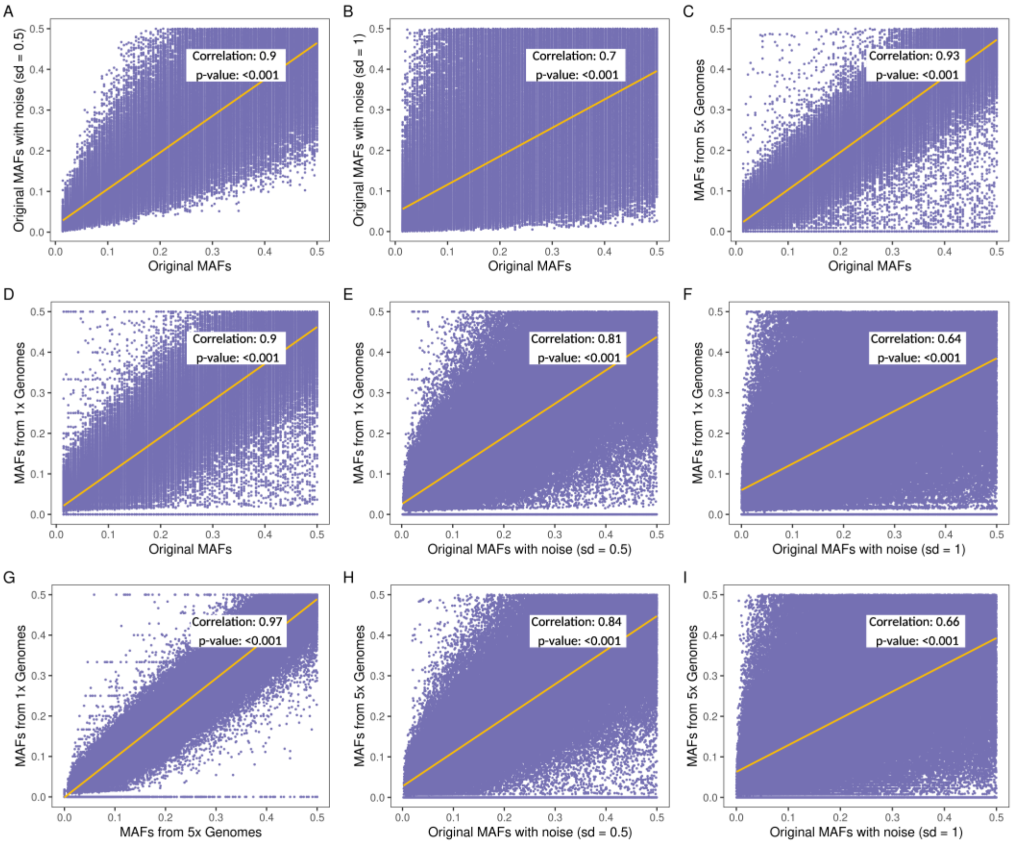


**Figure S3:** Minor allele frequencies (MAF) with different noise levels. “Original MAF” indicates TSI allele frequencies (perfect information), “Original MAF with noise (sd=0.5)” and “Original MAF with noise (sd=1)” indicate cases where Gaussian noise is added to the original TSI MAF. “MAF from 1x genomes” indicates MAF called using genomes of 1x coverage (n=72, comprising parent-offspring pairs), and MAF from 5x genomes” indicates MAF called using genomes of 5x coverage (n=72, comprising parent-offspring pairs) (Methods). The plots are drawn using 199,095 SNPs. The Spearman correlation coefficients and p-values are also shown.

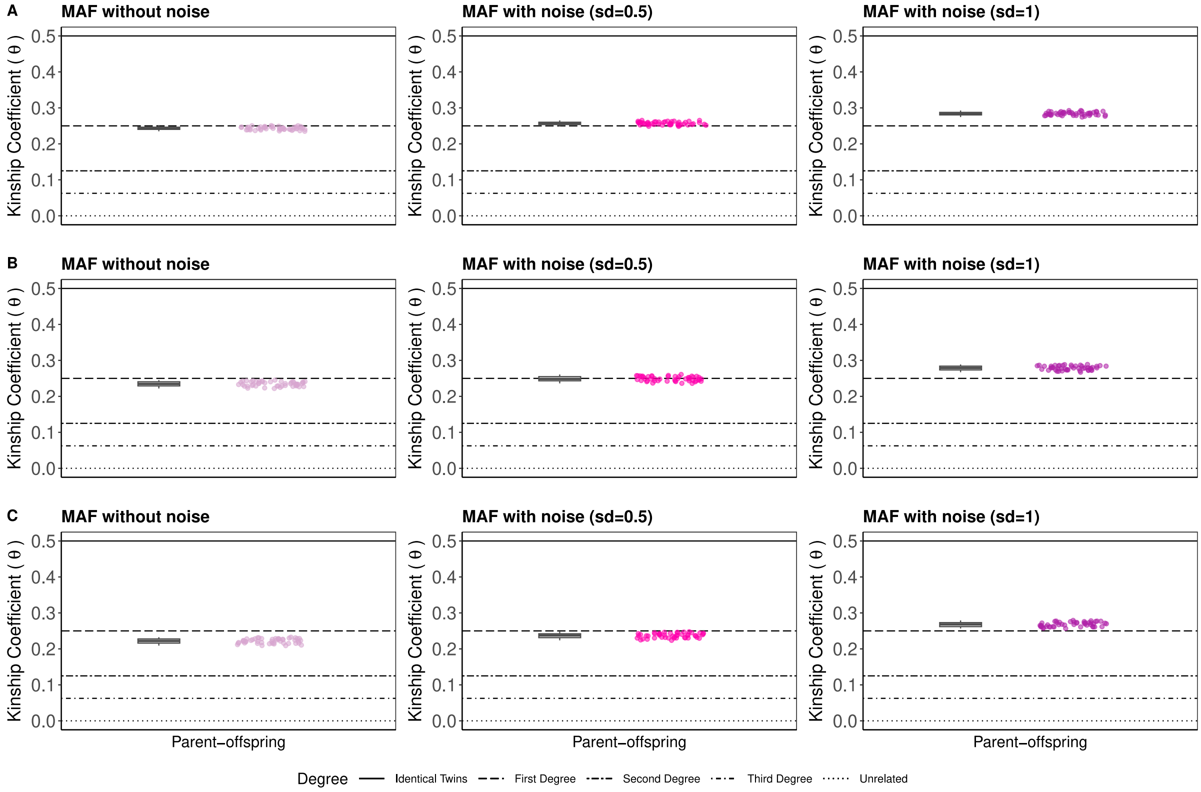


**Figure S4**: The effect of noise in $\theta$ estimates using NgsRelate on 48 parent-offspring pairs, without random noise or with different levels of random Gaussian noise added (sd=0.5 or =1). (A) $\theta$ estimates using the original MAF (i.e., perfect information). (B) $\theta$ estimates using MAF calculated using n=72 5x genomes. (C) $\theta$ estimates using MAF calculated using n=72 1x genomes. Horizontal lines show the theoretical $\theta$ values. The boxplots and jitter-added points show the distribution of the 48 points.

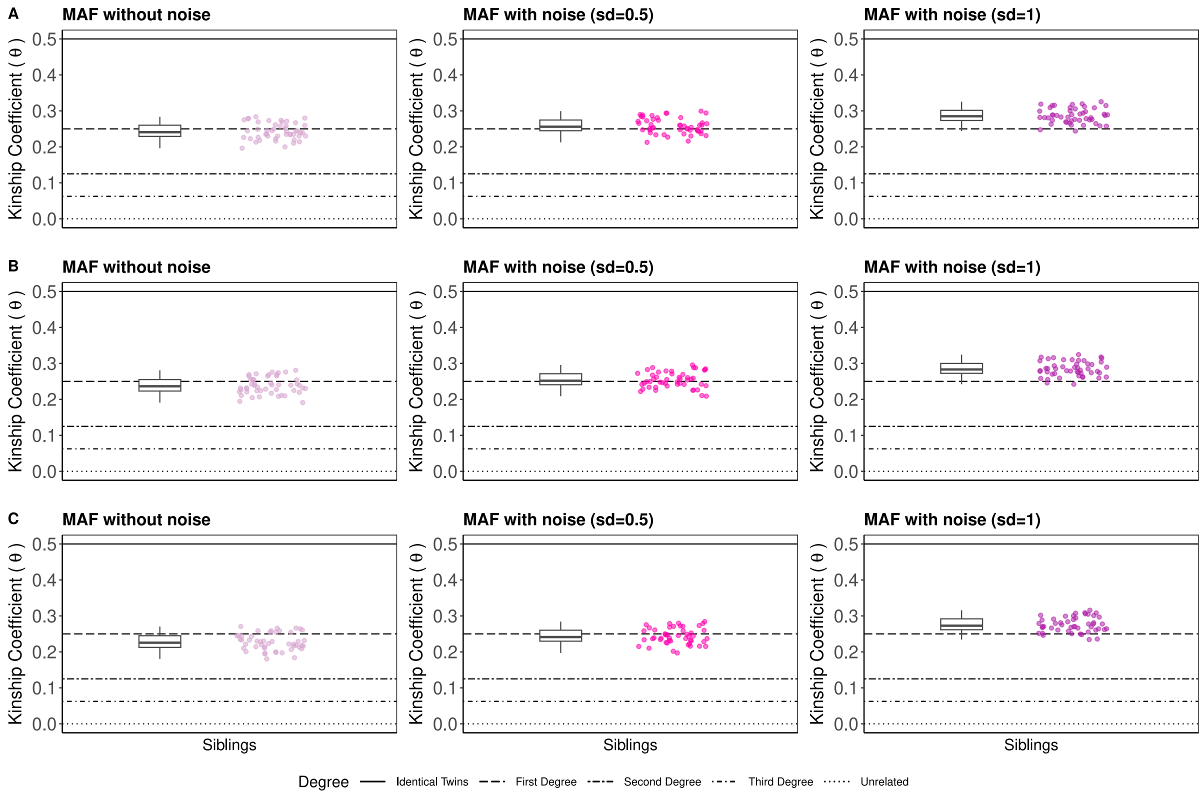


**Figure S5**: The effect of noise in $\theta$ estimates using NgsRelate on 48 sibling pairs, without random noise or with different levels of random Gaussian noise added (sd=0.5 or =1). (A) $\theta$ estimates using the original MAF (i.e., perfect information). (B) $\theta$ estimates using MAF calculated using n=96 5x genomes. (C) $\theta$ estimates using MAF calculated using n=96 1x genomes. Horizontal lines show the theoretical $\theta$ values. The boxplots and jitter-added points show the distribution of the 48 points.

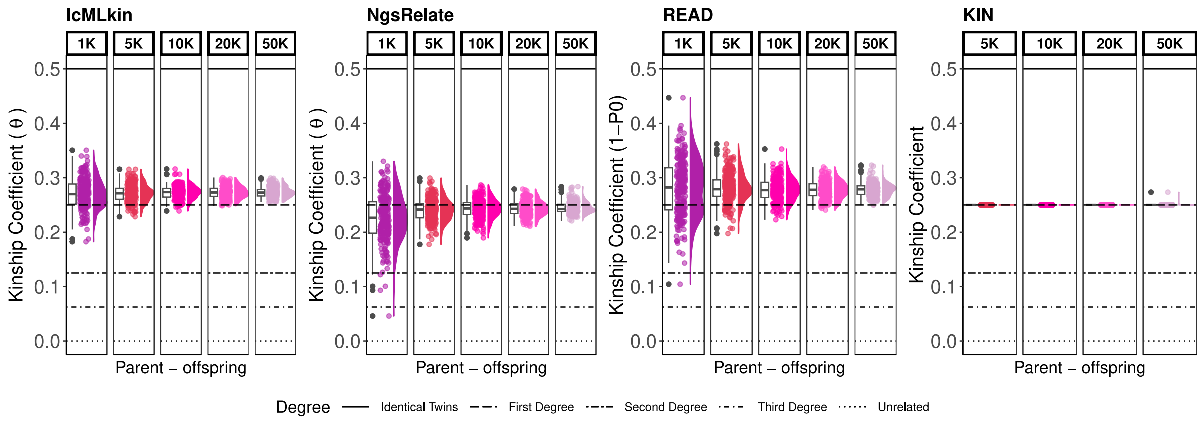


**Figure S6:** Kinship coefficient estimates for parent-offspring pairs in the presence of inbreeding by lcMLkin, NgsRelate, READ, and KIN. Horizontal lines show the theoretical kinship coefficients. The boxplots, jitter-added points, and density plots show the distribution of the same sample of 240 points. $\theta$ from NgsRelate was calculated ignoring the inbreeding-related Jacquard coefficients: $\hat{\theta} = J_7/2 + J_8/4$. KIN results for 1K are missing because the algorithm does not perform at this coverage.

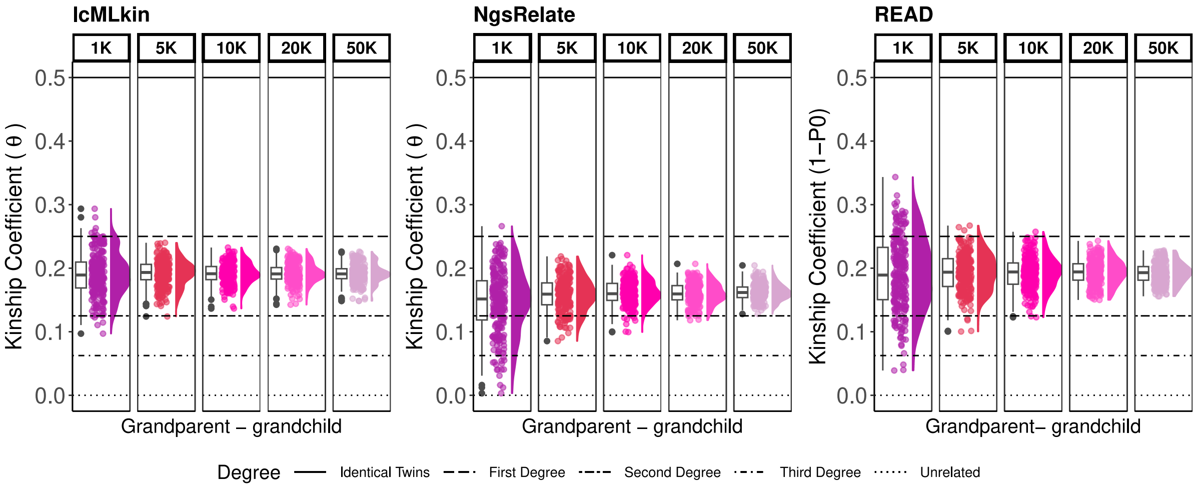


**Figure S7:** Kinship coefficient estimates for grandparent-grandchild pairs in the presence of inbreeding by lcMLkin, NgsRelate, READ, and KIN. Horizontal lines show the theoretical kinship coefficients. The boxplots, jitter-added points, and density plots show the distribution of the same sample of 240 points. $\theta$ from NgsRelate was calculated ignoring the inbreeding-related Jacquard coefficients: $\hat{\theta} = J_7/2 + J_8/4$. KIN results are missing because the algorithm did not perform at this relationship type.
